# Host-derived Reactive Nitrogen Species mediate the *Cryptococcus neoformans* yeast-to-titan switch via fungal-derived superoxide

**DOI:** 10.1101/2021.03.01.433276

**Authors:** Xin Zhou, Guillaume E. Desanti, Robin C. May, Ivy M. Dambuza, Elizabeth R. Ballou

**Affiliations:** Institute of Microbiology and Infection, School of Biosciences, University of Birmingham, Edgbaston, Birmingham, B15 2TT, UK; Medical Research Council Centre for Medical Mycology, University of Exeter, Exeter, EX4 4QD, UK

## Abstract

In the host lung, the human fungal pathogen *Cryptococcus neoformans* undergoes a morphological switch from small haploid yeast to large polyploid titan cell, contributing to *C. neoformans* virulence. Titan cells are less readily phagocytosed and can survive host nitrosative and oxidative stresses. We and others previously showed that titanization is triggered by host-relevant signals including CO_2_ and lung-resident bacteria, and addition of these factor is sufficient to induce titan cells *in vitro*. Here we investigate the molecular mechanisms that drive this transition and demonstrate that host-derived immune signals can increase the degree and frequency of titanization. Specifically, host-relevant reactive nitrogen species increase the accumulation of endogenous superoxide within cryptococcal cells, particularly within nuclei, where it can cause genotoxic stress. Consistent with this, we observe the accumulation of Rad51 protein, a marker of the double strand break repair pathway, in titanizing cultures. Blocking superoxide accumulation inhibits titanization, yet titanization also requires superoxide detoxification through Superoxide Dismutase (SOD) activity. Loss of mitochondrial Sod2 activity locks cells in the yeast phase, while Sod1 is required for the production of viable titan daughter cells. We hypothesize that the redox responsive transcription factor Yap1 in part mediates this response by regulating *SOD2/SOD1*. In addition, we show that Sod1 translocates to the nucleus, where it is likely involved in the detoxification of genotoxic superoxide. Together, these findings reveal a major new regulatory mechanism for the yeast-to-titan transition.

**Author Summary:** During fungal infection, host phagocytes produce reactive oxygen and nitrogen species (ROS/RNS), major determinants of infection outcome. Fungal pathogens have developed numerous strategies to neutralize and detoxify ROS, but RNS remain important effectors for infection control. In the lung, the human fungal pathogen *Cryptococcus neoformans* can undergo a morphological switch from small haploid yeast to large highly polyploid titan cells with increased ROS/RNS stress resistance, and the capacity to produce haploid or aneuploid daughters. Here, we report that RNS are a major signal driving the frequency and degree of titanization and act by increasing endogenous ROS within the fungus. We show that the accumulation of endogenous ROS is required for the yeast-to-titan transition, and is associated with increased genotoxic stress leading to polyploidy. Yet, failure to detoxify this ROS, either in mutants defective in Superoxide Dismutase activity or the oxidative stress response protein Yap1, impairs titan cell budding and reduces progeny viability. Therefore, the interface of exogenous RNS and endogenous ROS regulation during host-pathogen interaction represents an Achilles’ heel for this major human fungal pathogen.

## Introduction

*Cryptococcus neoformans* is a major human fungal pathogen that proliferates as a budding yeast and accounts for 15% of HIV-related deaths [1–3]. During infection of the lung, cryptococcal cells undergo an unusual morphological transition, changing from small haploid yeast to large, polyploid titan cells with increased oxidative/nitrosative stress and drug resistance [4–9]. We and others previously showed that the initial switch is triggered by host environmental conditions, including nutrient limitation, hypoxia, and bacterial peptidoglycan, abundant in the host lung [9–12]. However, the additional impact of host stimuli on fungal ploidy and morphology remains unclear. Here we show that host factors generated by the immune system can affect fungal titanization by triggering the endogenous generation of an important signalling molecule, the superoxide radical, within fungal cells.

In the host lung, *C. neoformans* yeast proliferate both extracellularly and within phagocytes [13, 14]. Successful control of infection requires a robust adaptive response, including T-cell mediated activation of phagocytic macrophages, which bind and internalize fungal cells [15, 16]. Activated macrophages are polarized (M1/M2) and this polarization is dynamic, with an early M2 profile shifting to a protective M1 profile over the course of infection [16, 17]. Among other responses, M1 cells express Nox2, which generates superoxide anion (O_2_^-^) and other oxygen-derived intermediates, and inducible nitric oxide synthase (iNOS), which generates nitric oxide and its derivatives [18]. These host-derived reactive oxygen and reactive nitrogen species (ROS/RNS) are well recognized for their important roles in controlling pathogens by causing extensive damage to DNA, proteins and lipids [19, 20]. For *C. neoformans* in particular, host-derived RNS, together with ROS, control fungal proliferation and dissemination [14].

In response to host oxidative and nitrosative stresses, *C. neoformans,* like other pathogenic fungi, has evolved stress resistance responses that enable fungal survival and lead to disease [21–24]. Typical fungal responses involve the expression of ROS/RNS detoxification enzymes including superoxide dismutase (SOD), catalase (CAT), glutathione peroxidases, thioredoxin, and peroxiredoxins [19, 24–27]. These enzymes are required for fungal cells to detoxify intracellular ROS or RNS and play important roles in fungal biology and pathogenicity. For example, in *C. neoformans*, *Candida albicans*, and *Aspergillus fumigatus,* the loss of SOD activity has been shown to have direct consequences for fungal virulence [28–31]. *C. neoformans* additionally detoxifies host ROS through the expression of polysaccharide capsule and melanin [32].

In addition to exogenous ROS/RNS, mitochondria are a major source of endogenous ROS within the fungal cell. During phagocytosis, it is predicted that NO^-^, the main product of the iNOS pathway, targets fungal mitochondria leading to the accumulation of Complex III-derived O_2_^-^ within the fungal cell [33]. Consistent with this, host-derived RNS blocks *C. neoformans* proliferation, and iNOS activity is required for host survival [34, 35]. Detoxification of RNS via nitric oxide dioxygenase and S-glutathione dehydrogenase as well as nitrosative stress responsive transcription factors are required for *C. neoformans* virulence [27, 36, 37].

Endogenous ROS also act as signalling molecules regulating different biological and physiological processes [24, 38–43]. In *A. nidulans* and *C. albicans*, the endogenous generation of a superoxide gradient was shown to regulate polarised growth and sexual development, and filamentation, respectively [39–41]. A similar role for endogenous superoxide has not yet been described in *C. neoformans*.

Here, we examined the *in vitro* interaction of host phagocytes with pre-titanized cells [11, 44]. Upon co-culture, we observed a significant further increase in cryptococcal cell size and an increase in endogenous superoxide, the generation of which is required for titanization. Through mutant analysis and live cell imaging we begin to dissect the molecular mechanisms governing this process. We demonstrate that, rather than simply killing fungal cells or inhibiting fungal growth, host phagocyte iNOS activity can augment the formation of large titan cells, with implications for host survival.

## Results

To study the yeast-to-titan transition, fungal cells can be induced to form titan cells using our established protocol: growth in minimal media followed by exposure for 24 hr to 10% Fetal Calf Serum (FCS), a source of bacterial peptidoglycan [9, 11]. This yields a mixed culture of cells proliferating either via titan (15-20%, >10 μm) or yeast (<10 μm) growth patterns, and throughout this manuscript we define cultures primed in this way as “pre-titanized”. To study how this transition is impacted by interaction with the host, we performed time-lapse imaging of pre-titanized cells subsequently co-cultured with murine J774.1 macrophage cell line stimulated with interferon gamma (IFN-γ) (to simulate M1-like conditions) (Fig 1A and S1A Fig). We observed that engulfed H99 titan cells continue to undergo cell enlargement within the phagolysosome. In addition, we noticed an overall increase in the maximum size and frequency of titan cells following co-culture relative to growth in pre-titanizing conditions [9, 11]. This increase was not observed for control cells that had not been primed by previous exposure to FCS were co-incubated with phagocytes, indicating that host phagocytosis alone is not sufficient to trigger titanization (S1B Fig). This striking observation raised the possibility that, in addition to a role for nutrient limitation and bacterial peptidoglycan, host factors may also impact the degree and frequency of the yeast-to-titan switch in the context of the host. We therefore set out to understand the molecular mechanisms driving this fungal response to the host environment.

**Figure 1:**
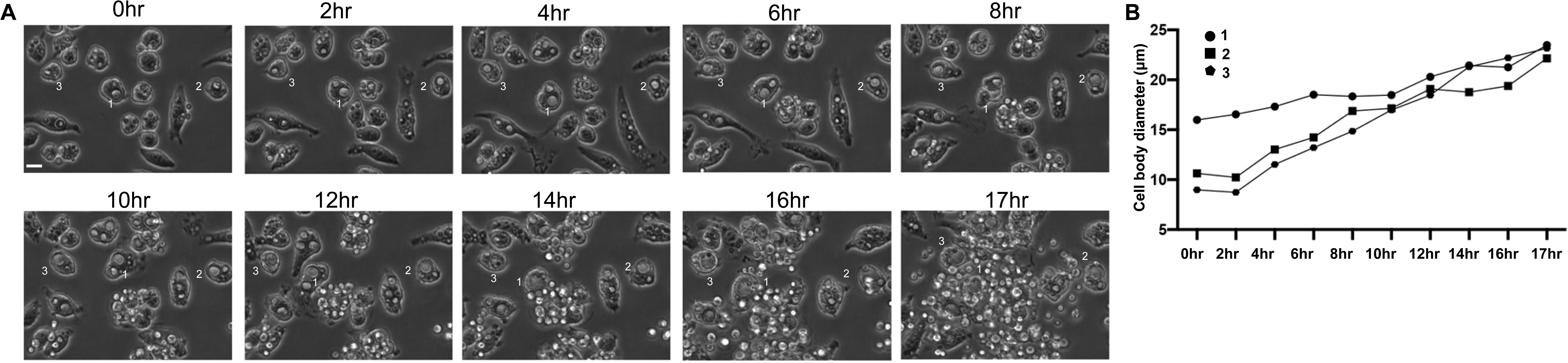
Time-lapse images from co-incubation of pre-induced H99 with macrophages. *C. neoformans* cells were incubated under titan inducing conditions for 24hr then co-incubated with J774 murine macrophages. Co-cultured cells were imaged for 18hr. (A) Representative micrographs are shown at 2 hr intervals; Indicate cells are representative of engulfed cryptococcal cells that increased in size (B) Quantification of cell diameter for indicated cells over time.

## Host-derived immune signals induce formation of large titan cells

We first asked whether activation of host immune cells is required to amplify fungal cell size. We used murine bone marrow derived macrophages (BMDMs) isolated from C57BL/6 (WT) to repeat the co-incubation assays. BMDMs are professional phagocytes that can be primed to be either M1 or M2, where M1 cells are strong producers of NO^-^, TNF, and IFN-γ, factors important for the control and clearance of cryptococcal cells [45, 46]. Treatment with LPS and IFN-γ stimulates M1 polarization, which is associated with macrophage-mediated fungal killing via increased phagosomal ROS and NO^-^ [47]. In contrast, treatment with IL-4 stimulates an M2 polarization associated with macrophage mediated clearance.

As shown in Fig 2A, cryptococcal cells co-incubated with basally activated murine BMDMs increased in size relative to fungal cells alone, similar to co-incubation with J774.1 macrophages (Fig 2A). Of the engulfed fungal cells, approximately 50% increased further in cell size, while the remainder either proliferated as yeast or were controled. We then activated BMDMs with either LPS, IFN-, or both LPS and IFN-γ, priming M1 polarization, and examined fungal cell size by live cell imaging for 18 hr (Fig 2A). There was a significant increase in fungal cell size when co-incubated with unstimulated BMDMs compared to growth in matched medium alone (p<0.0001). Stimulating BMDMs with either LPS or IFN-γ or with both LPS and IFN-γ further increased fungal cell size compared to fungal cells in matched media (Fig 2A, p<0.0001). Representative images are shown in Figure 2B. There was no statistically significant difference between fungal cell populations incubated with either BMDM alone or BMDMs stimulated with LPS and IFN-γ, however we consistently observed an increase in frequency of very large titan cells (>15 μm) in co-stimulated cultures (Fig 2A). Flow cytometry of fungal cells after lysis and gating to exclude BMDMs confirmed the observed increase in fungal cell size, with the most dramatic size increase occurring after dual activation (FSC-A) (Fig 2C, gating strategy S2A Fig).

**Figure 2:**
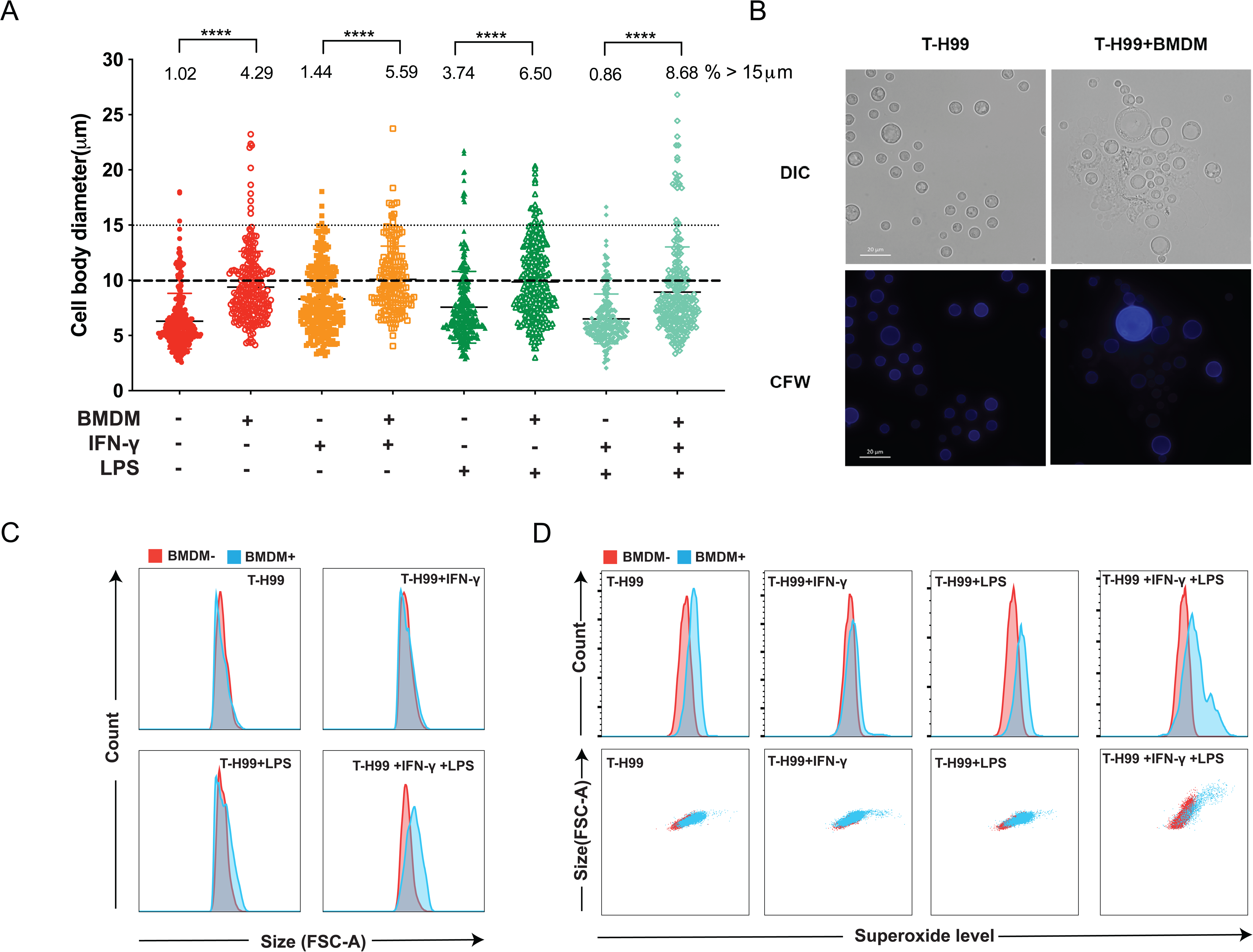
Fungal cell size increases after co-incubation with BMDMs and is influenced by BMDM activation. (A) *C. neoformans* pre-titanized cell size changes after different incubation assays: *C. neoformans* were grown in RPMI in presence or absence of BMDMs, IFN-γ and/or LPS. *C. neoformans* cell size was assessed for individual cells (cell body diameter, excluding capsule; n>200). Data were assessed for normality by Shapiro-Wilk and analysed by one-way ANOVA, \*\*\*\**p*<0.0001. (B) Microscopy images of cells grown in RPMI (*left*), or after co-incubation with BMDMs activated with IFN-γ and LPS (*right*, BMDMs lysed with cold sdH_2_O); Cells were stained with calcofluor white (CFW) for cell wall (*blue*). Scale bar: 20 μm. (C,D) Flow cytometric analysis of cell size (C) and superoxide staining distribution (D) for *C. neoformans* cells co-incubated with activated BMDMs. Fungal cells were identified using gates for capsule (S2A Fig). Single cells were identified by FSC-A vs FSC-H to remove doublets. D) Histograms for superoxide levels for fungal cells from indicated conditions (*top panel*); size (FSC-A) vs superoxide level scatter plots for cells from indicated conditions (*bottom panel*). Three biological replicates were performed and a single representative replicate is presented. “T” indicates pre-titanized.

We asked whether stimulation with IL-4 impacted the capacity of BMDMs to increase fungal cell size. Fungal cell size was determined by flow cytometry following gating to exclude BMDMs as above. In contrast to stimulation with LPS and IFN-γ treatment with IL-4 reduced the proportion of very large titan cells upon co-incubation with BMDMs (S2B Fig). Together, these data suggest that exposure to stimulated M1, but not M2, immune cells, can induce the formation of large titan cells.

M1-polarized phagocytes are strong producers of phagosomal O_2_ ^-^and NO^-^, factors important for control and clearance of *Cryptococcus* cells [14, 47]. Intriguingly, we observed that superoxide levels within cryptococcal cells increased in parallel with cell size after co-incubation with BMDMs. As shown in Fig 2D, cryptococcal cells exposed to BMDMs showed higher levels of superoxide compared with *in vitro* titanized cells alone, and the most dramatic increase in superoxide was observed for co-stimulated cells (Fig 2D top panel). The shift in superoxide correlated with increased cell size for fungal cells co-incubated with BMDMs for all conditions compared to fungal cells exposed to LPS and/or IFN-γ in the absence of BMDMs, further demonstrating that this effect was mediated by BMDM activity (Fig 2D).

## Exogenous nitrosative stress is sufficient to further potentiate titanization

The observation that co-incubation with M1 but not M2 macrophages concomitantly increased superoxide and cell size raised the hypothesis that host-derived ROS/RNS influence fungal morphogenesis. To investigate the roles of host-derived ROS and RNS in titanization, we directly added exogenous reactive oxygen (H_2_O_2_) or nitrogen (NO^-^) species donors (Glucose oxidase or DPTA NONOate) at levels consistent with the host environment to titan cultures at the time of induction[40, 48]. Treatment with exogenous H_2_O_2_ was associated with increased superoxide staining within fungi, suggesting that the cells are under ROS stress, and a moderate shift in median cell size was observed (Fig 3A). In contrast, the presence of exogenous NO^-^ stimulated increased O_2_ and nearly doubled median cell size (Fig 3A). These data suggest that host-derived RNS, and to a lesser extent ROS, can further potentiate titanization.

**Figure 3:**
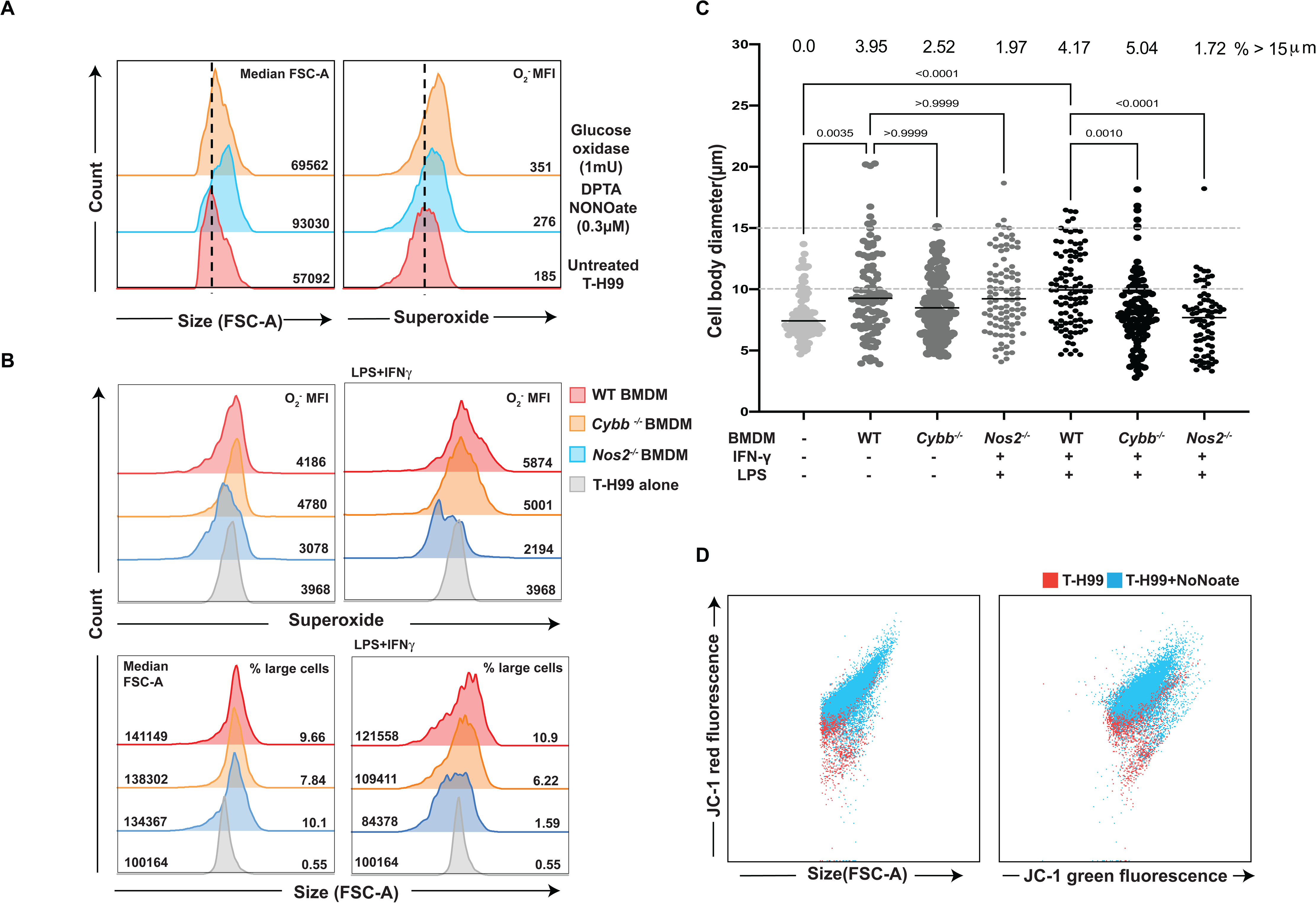
Exogenous RNS treatment correlates with increases in both cell size and superoxide level. (A) Histograms of size (FSC-A) and superoxide levels of titanized cells from untreated (*bottom*), DPTA NONOate (0.3μM) treated (*middle*), and Glucose oxidase (1mU) treated (*top*), Dotted lines indicate median fluorescence intensity (MFI) of untreated cells. (B) *C. neoformans* pre-titanized cell size changes after different incubation assays: *C. neoformans* were grown in RPMI in presence or absence of WT or *Cybb*^-/-^ or *Nos2*^-/-^ BMDMs activated or not with IFN-γ and LPS. Cell size was assessed for individual cells (diameter) for the cell body (capsule excluded; n>100). Data were assessed for normality by Shapiro-Wilk and analysed by one-way ANOVA, \*\*\*\**p*<0.0001, \*\*\**p*=0.0010. (C) Flow cytometer results of cell sizes and superoxide levels: *C. neoformans* cells were gated by capsule staining, then checked for superoxide level and size (FSC-A). Histograms of superoxide levels (*top panel*) and size (FSC-A) (*bottom panel)* for cells from different culture conditions. (D) Left: Size (FSC-A) vs JC-1 red fluorescence scatter plots of titanized cells from untreated (*red dots*) and NoNoate treated (*blue dots*) cultures; right: JC-1 green fluorescence vs red fluorescence scatter plots of titanized cells from untreated (*red dots*) and NoNoate treated (*blue dots*) cultures. Cells were stained with superoxide detector or JC-1, then analysed by flow cytometer. Single cells were identified by FSC-A vs FSC-H to remove doublets. Three biological replicates were performed and a single representative replicate is presented. “T” indicates pre-titanized.

To determine the physiological relevance of the roles of exogenous ROS vs. RNS, we co-incubated pre-titanized fungal cells with BMDMs from mice lacking either NOX (O_2_^-^, *Cybb*^*-/-*^) or iNOS (NO^-^, *Nos2*^*-/-*^) activity (Fig 3B, C). Fungal cell size was examined both by flow cytometry (Fig 3B) and microscopy (Fig 3C). In the absence of LPS and IFN-γ stimulation, there was no difference in cell size distribution between wildtype BMDMs and BMDMs from iNOS-deficient mice (Fig 3B). *Cybb*^-/-^ BMDMs lacking the Gp91^PHOX^ protein required for O_2_^-^ production were moderately less able to induce titan cell size increases compared to *Cybb*^+/+^ BMDMs (Fig 3B, C), but induced comparable endogenous superoxide. This suggests that host-derived ROS are minor contributors to fungal cell size increase [49, 50]. In contrast, when pre-titanized fungal cells were incubated with stimulated BMDMs lacking *NOS2* (iNOS-), there was a profound defect in fungal cell size increase relative to WT BMDMs (Fig 3B, C). In addition, endogenous superoxide was markedly lower in fungal cells co-incubated with iNOS-deficient mice (Fig 3B). Together, these data confirm that host-derived RNS are the major contributor to observed increases in fungal cell size.

## Exogenous RNS increases mitochondrial potential

Exogenous and endogenous ROS/RNS can cause dysfunction of mitochondria leading to cell death, however moderate exposure to exogenous RNS has been shown to increase cell respiration and endogenous ROS formation [19, 51–54]. While the impact of exogenous ROS and RNS on *C. neoformans* stress responses has been studied at the phenotypic, transcriptional and translational levels, their specific impact on mitochondrial function and morphology remains unclear [26, 27, 30, 55, 56]. In order to determine whether exogenous RNS impacts *C. neoformans* mitochondrial function, we measured mitochondrial membrane potential, an indicator of mitochondrial oxidative phosphorylation, during exposure to physiological concentrations of nitric oxide[48]. We used the fluorescent probe JC-1, which exists as monomer and emits green fluorescence at low membrane potential. In mitochondria with high membrane potential, JC-1 forms aggregates and emits orange-red fluorescence (S3 Fig). As shown in Fig 3D, treatment with NONOate increased the mitochondrial potential of pre-titanized cells, with a clear increase in JC-1 red fluorescence relative to untreated pre-titanized cells. This suggests that exogenous RNS increases mitochondrial potential and mitochondrial ROS production.

## Endogenous superoxide levels are dynamic throughout titanization and correlate with the capacity of isolates to form titan cells

Together, our data suggest that host-derived RNS, and to a lesser extent ROS, influence endogenous fungal superoxide, resulting in fungal size increase. We hypothesised that endogenous fungal superoxide influences the changes in fungal cell size observed during the 24hr yeast-to-titan switch, even in the absence of host stimulation. We therefore characterised endogenous superoxide levels over the course of titanization. We monitored endogenous superoxide using a superoxide-specific dye from 4hr post-induction, when titan cells are not apparent, until 24hr, when titan cells comprise 15% of the population. Flow cytometry showed that endogenous superoxide levels are initially high, then decrease 8hr post-induction, becoming bi-phasic (Fig 4A). From 13 to 18 hr, there is an overall shift to the right, and a minority of cells generate high levels of superoxide (Fig 4A). This minority population is maintained over the subsequent 12 hr, at which point overall superoxide is stabilized at an intermediate level. After 24 hr, the subset of cells with very high ROS correlate with the largest cells in this population (Fig 4A). When we monitored mitochondrial activity over time using the JC-1 probe, we observed a similarly high initial potential (red fluorescence) that decreased by 8 hr and then stabilized and was maintained at a high level from 13 hr (Fig 4B). We conclude that during titanization, endogenous superoxide levels are dynamic across the population and are reflective of high mitochondrial activity.

**Figure 4:**
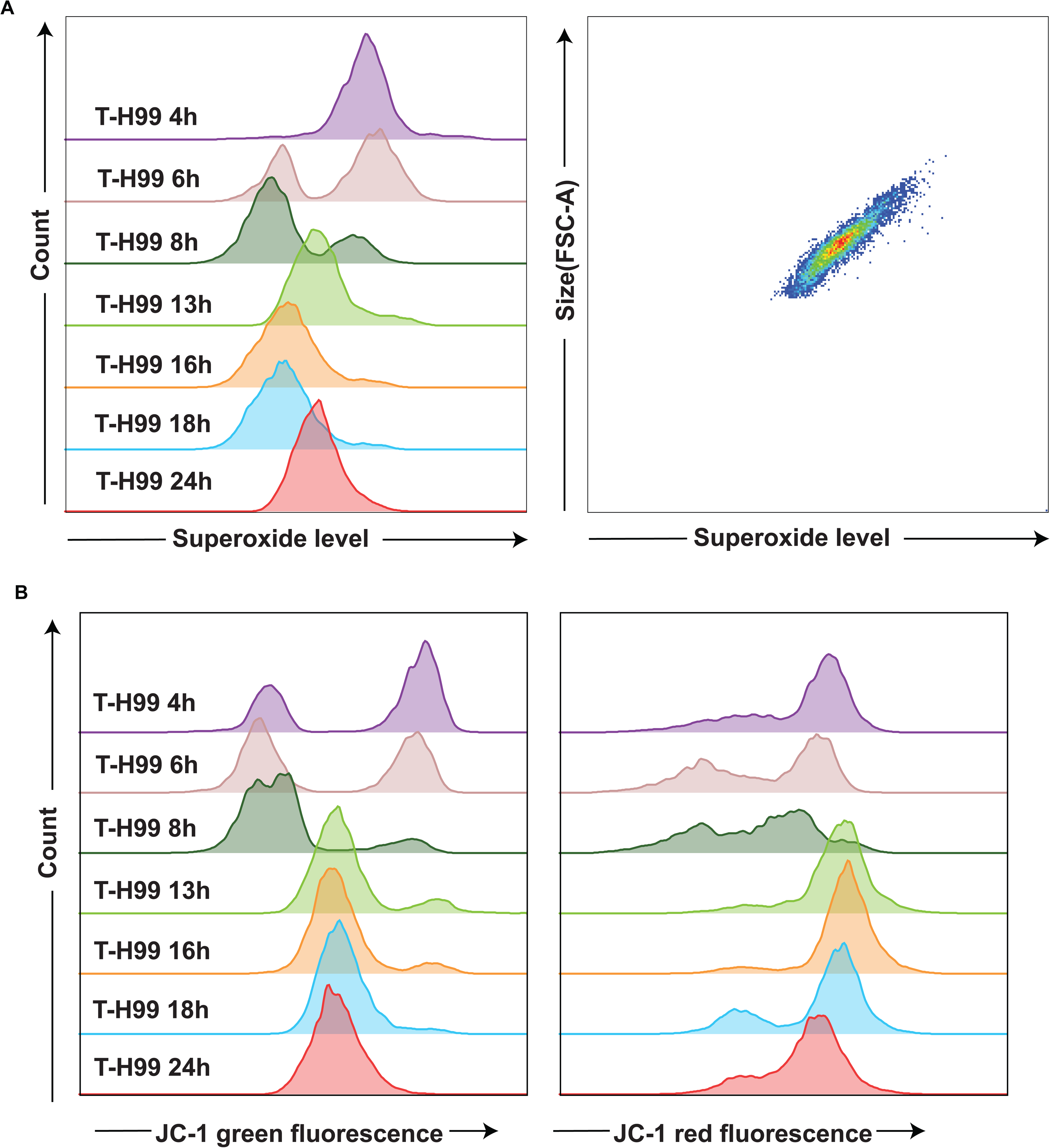
Superoxide accumulation is dynamic during titanization. (A) Histograms of superoxide levels of *C. neoformans* cells from different time points over titanization from 4hr to 24hr post-induction. size (FSC-A) vs superoxide level scatter plots for cells from 24hr post-induction. Cells were stained with superoxide detector and analysed by flow cytometer. (B) Histograms of JC-1 green fluorescence (*left*) and JC-1 red fluorescence (*right*) of *C. neoformans* cells from different time points over the yeast to titan transition from 4hr to 24hr post-induction. Cells were stained with superoxide detector or JC-1, then analysed by flow cytometer. Single cells were identified by FSC-A vs FSC-H to remove doublets. Three biological replicates were performed and a single representative replicate is presented. “T” indicates pre-titanized.

The dynamic nature of superoxide levels suggests that ROS detoxification is also important to fungal morphogenesis. Consistent with this, titan-induced H99 cells were more sensitive to killing by exogenous H_2_O_2_ than YNB-grown H99, suggesting titanized cultures are under increased endogenous oxidative stress (Figure 5A). Moreover, we observed a correlation between the capacity of clinical isolates to undergo titanization and sensitivity to exogenous H_2_O_2_ when grown under titan-inducing conditions: The clinical isolate Zc8, like H99, forms titans and becomes sensitive to H_2_O_2_ upon titan induction. In contrast, after exposure to titan conditions the Zc1 hypo-titanizing isolate and Zc12, which does not form titans, are no more sensitive than uninduced cells (Fig 5A). Microscopy and flow cytometry to measure superoxide levels in induced cells revealed that, after *in vitro* titan induction, H99 and Zc8 exhibit higher endogenous superoxide levels compared to non-titanizing isolates (Fig 5B, C). Co-incubation of the clinical isolates with human blood monocytes revealed that the impact of phagocytes on fungal cell size also correlates with the capacity to titanize. After 24hr co-incubation with monocytes, H99 and Zc8 increased in maximum cell size up to >25 μm (H99) and >15μm (Zc8) (Fig 5D), sizes which were only detected *in vivo* previously [44]. In contrast, Zc1 and Zc12 did not exceed the 10 μm threshold even after co-incubation with human monocytes (Fig 5D).

**Figure 5:**
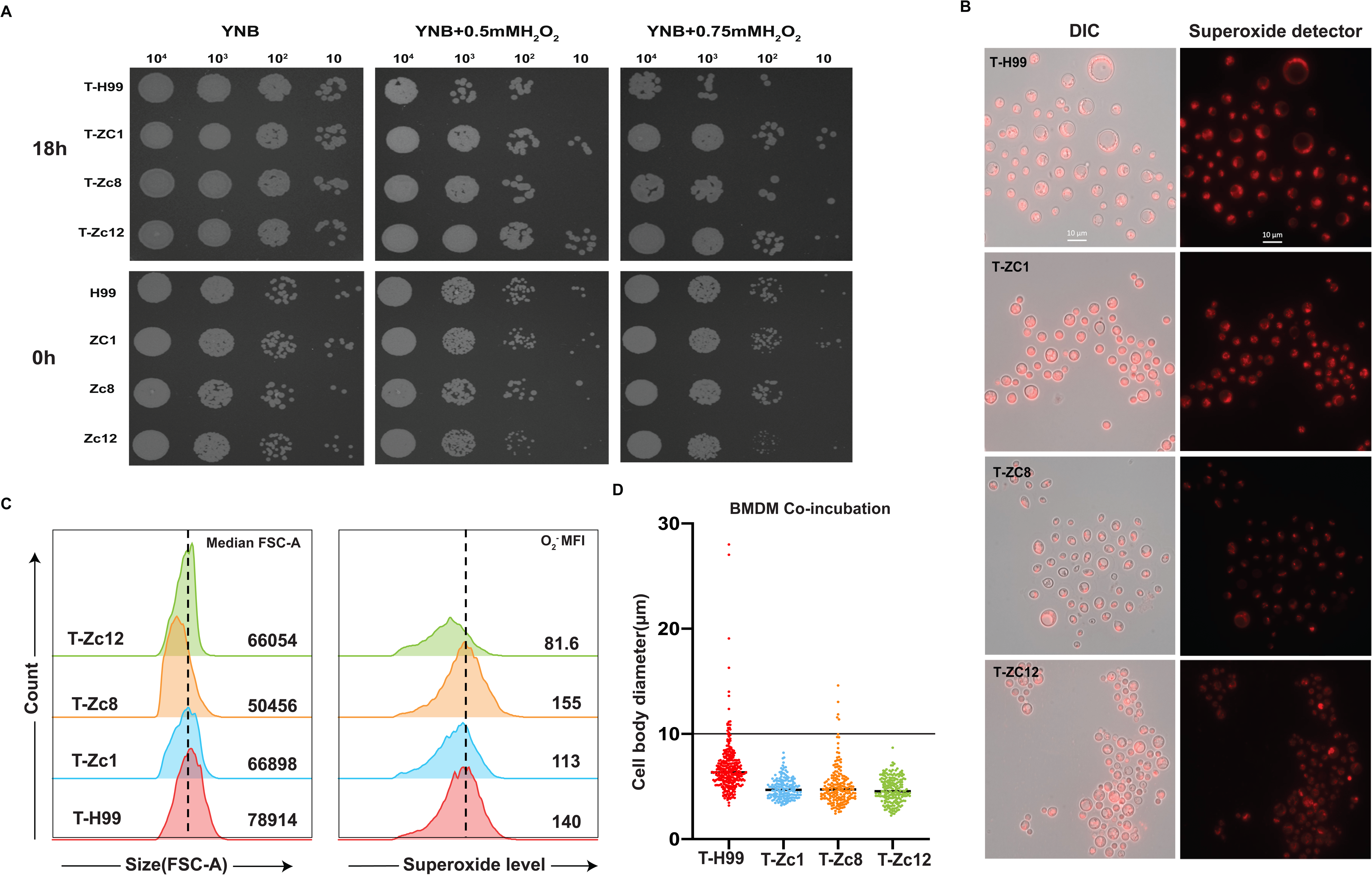
Superoxide increase with sizes within clinical isolates after co-incubation with monocytes. (A) Hydrogen peroxide stress spot plates of 3 clinical isolates (Zc1, Zc8, Zc12) with H99 as a control: cells were pre-grown in YNB and directly spotted onto plates (0hr) or incubated in 10% FCS in PBS, 5%CO2, 37°C for 18h(18h), and then plated on YPD plates with or without H_2_O_2_. (B) Micrographs of cells stained with superoxide detector. Scale bar: 20μ superoxide levels among indicated strains. Cells were stained with superoxide detector and analysed by flow cytometer. Single cells were identified by FSC-A vs FSC-H to remove doublets. Dotted lines indicate MFI of H99 titanized cells. (D) Cell sizes of H99 and 3 clinical isolates after *in vitro* induction (24hr) and co-incubation with monocytes: Cell size was assessed for individual cells (cell body diameter, excluding capsule; n>200). Three biological replicates were performed and a single representative replicate is presented. “T” indicates pre-titanized.

Finally, we visualized endogenous ROS in fungal cells after co-culture with monocytes: superoxide was apparent as bright cytoplasmic foci and in the nucleus in H99 and Zc8 titan cells, yet was localised to diffuse puncta in H99 and Zc8 yeasts and titanides. Similar diffuse puncta were observed in all cells in the non-titanizing Zc1 and Zc12 isolates (Fig 5B). Overall, these data suggest that both level and localization of superoxide dictate phenotypic outcome and that this occurs regardless of genetic background.

## Endogenous superoxide is required for titan cell formation yet must be detoxified for the generation of large cells

To determine whether superoxide is required for titan cell formation, we treated H99 titan cultures with MnTBAP, a SOD mimetic drug. MnTBAP is readily taken up by live cells and can convert endogenous superoxide into H_2_O_2_, which is subsequently detoxified by fungal catalases [56–58]. Treatment with MnTBAP blocked titan cell formation, with no cell larger than 10 μm or with ploidy >8C observed (Fig 6A, B). Flow cytometry confirmed that treatment decreased superoxide accumulation (Fig 6C). Direct exposure to exogenous H_2_O_2_ did not affect titanization (Fig 3A), suggesting that the MnTBAP-mediated block in titanization is not the result of increased H_2_O_2_ stress. No defect in growth was observed, and treated cells resembled “typical” yeast cells (<10 μm) in morphology (S4 Fig).

**Figure 6:**
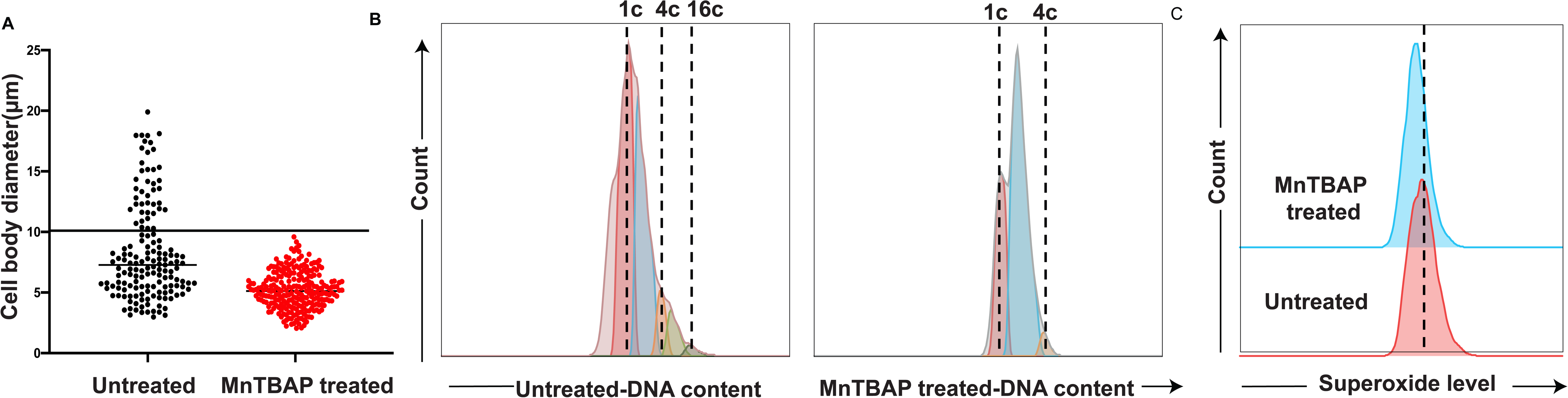
Titan cell formation can be inhibited by reducing the accumulation of superoxide. (A) Cell body diameter for *C. neoformans* H99 cells induced to form titans for 24hr in the absence or presence of the SOD mimetic drug MnTBAP (1mM). Cell size was assessed for individual cells (cell body diameter, excluding capsule; n>200). (B) DNA content (DAPI) of untreated (*left*) and MnTBAP treated (*right*) cells was assessed by flow cytometer. YPD grown yeast cells from matched parent strains and MFI 1C and 2C populations were used to identify 1C and 2C peaks (Fig S3B). The dotted lines indicate MFI of different DNA content peaks. (C) Histograms of superoxide levels of titanized cells from untreated and MnTBAP treated cells. Cells were stained with superoxide detector and analysed by flow cytometer. (B&C) Single cells were identified by FSC-A vs FSC-H to remove doublets. Dotted lines indicate MFI of untreated cells. Three biological replicates were performed and a single representative replicate is presented.

These data pose an important biological challenge for the fungal cell: the generation of superoxide is a requirement for titanization, and host factors that increase superoxide also increase fungal cell size, yet excessive endogenous ROS can cause oxidative damage and sensitizes the fungus to exogenous ROS stress and host cell killing. Balancing this response requires the activity of the oxidative stress response pathway.

The AP-1-like transcription factor Yap1 is a stress response protein induced by oxidative stress in *C. neoformans,* and loss of *YAP1* sensitizes cells to a wide variety of oxidative stress compounds (H_2_O_2_, t-BOOH (H_2_O_2_), diamide (O_2_^-^)) [59, 60]. Consistent with a defect in ROS detoxification, examination of *yap1Δ* titan-induced cultures revealed two key changes (Figure 7A, B): 1) we did not detect highly polyploid cells in the *yap1Δ* mutant (>16C) and 2) there was a reduction in the percentage of titanides, cells smaller than 5 μm with “sub-1C” DNA content that originate from titan cells [44]. We hypothesized that this might be due to failure of *yap1Δ* cells to detoxify high endogenous superoxide levels observed over the course of titanization (Fig 4A). Under basal conditions (YPD, 30°C), Yap1 regulates oxidative resistance genes such as *SOD1* and *SOD2,* which detoxify superoxide to H_2_O_2_, which is then converted to O_2_ and H_2_O by catalases (*CAT*1-4) [59, 60].

**Figure 7:**
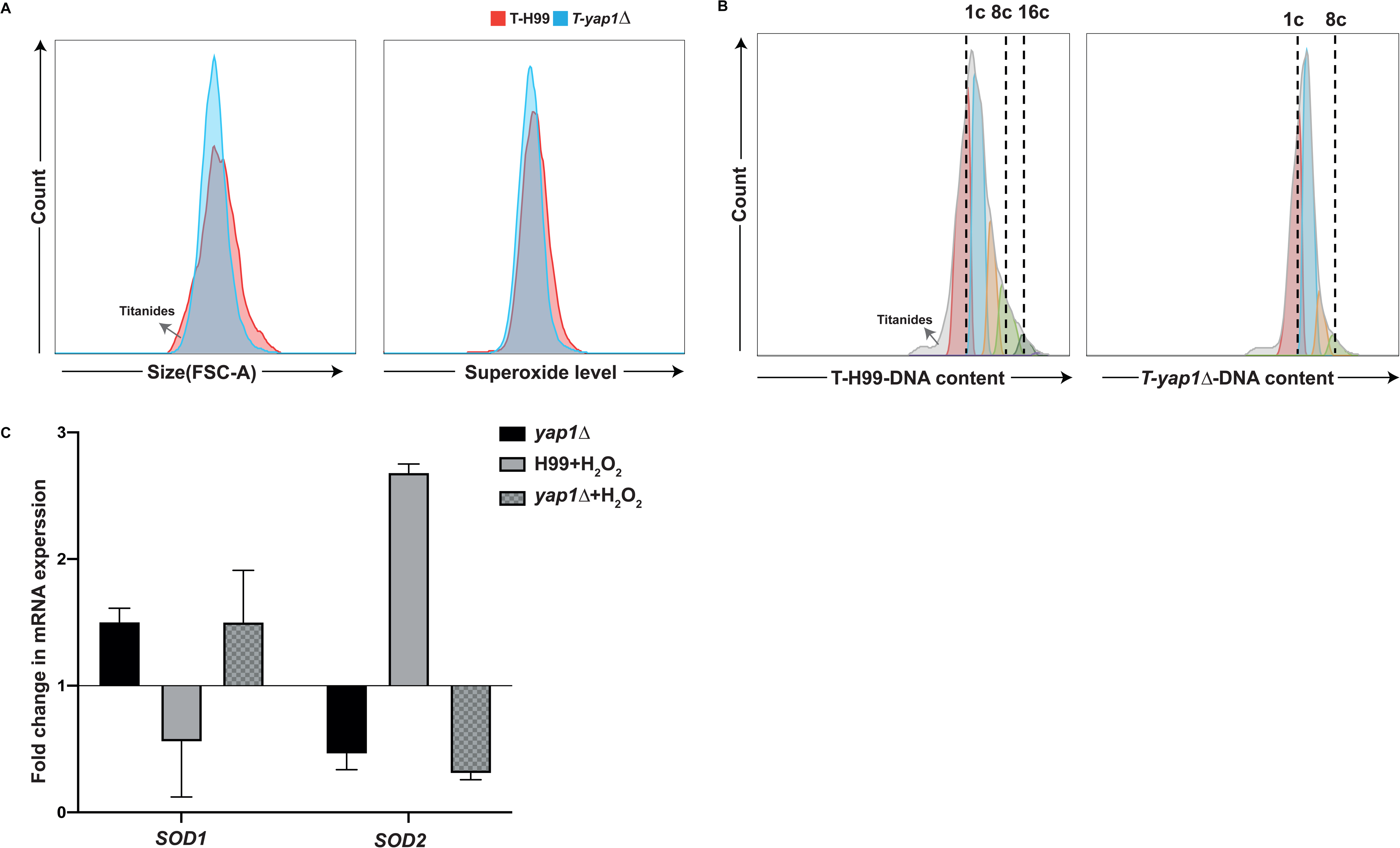
Yap1 regulates ROS detoxification and the degree of titan cell formation. (A) Histograms of cell size and superoxide levels of T-H99 (*red*) and T-*yap1Δ* (*blue*) after 24hr titan induction. Cells were stained with superoxide detector and analysed by flow cytometer. (B) DNA content of titan induced H99 and *yap1Δ* cells (24hr). YPD-grown yeast cells from matched parent strains and MFI 1C and 2C populations were used to identify 1C and 2C peaks. The dotted lines indicate MFI of different DNA content peaks. (A&B) Grey arrows indicate titanide cells. Single cells were identified by FSC-A vs FSC-H to remove doublets. (C) Relative mRNA expression levels of *SOD1* and *SOD2* in H99 treated with1.5mM H_2_O_2_ relative to untreated H99 and *yap1Δ*with or without 1.5mM H_2_O_2_ relative to untreated H99. “T” indicates pre-titanized.

However, data are lacking about the role of Yap1 under oxidative stress. We asked whether *SOD* regulation is altered upon mild oxidative stress (YPD + 1.5 mM H_2_O_2_). This experimental condition was selected instead of titan inducing conditions because the heterogeneous nature of titan cultures complicates the interpretation of transcriptional data during the early phases of induction. Instead, we compared the expression of these two genes in H99 and *yap1Δ* during yeast-phase growth (YPD, 30 deg, +/-1.5mM H_2_O_2_ for 5 hr) relative to untreated H99 via RT-PCR. As shown in Figure 7C, in the *yap1Δ* mutant under basal conditions, *SOD1* is induced relative to wild-type while *SOD2* expression is reduced relative to wild-type levels. Upon incubation with moderate concentration of H_2_O_2_ (1.5mM) to activate Yap1, wild-type expression of *SOD1* decreases compared with untreated cells, while *SOD2* exhibits a more than 2-fold increase (Fig 7C). In *yap1Δ*, H_2_O_2_-induced regulation of *SOD1* and *SOD2* is lost: both *SOD1* and *SOD2* behave similar to untreated *YAP1* cells (Fig 7C). We hypothesize that this loss of Superoxide Dismutase activity impairs titanization in the *yap1Δ* mutant.

## SOD2 is required for the initial detoxification of superoxide

To directly test the hypothesis that ROS detoxification influences titanization, we examined the requirement for components of this pathway including superoxide dismutase (*SOD1, SOD2*) and catalase (*CAT1, CAT2, CAT3, CAT4*). Deletion mutants were assessed for cell size and endogenous superoxide levels after exposure to inducing conditions for 24 hr. As shown in Fig 8A, neither *sod1, cat1, *Δ Δ*cat2* Δ*, cat3*Δ*, cat4* Δ single mutants, nor the *cat1/2/3/4* Δ quadruple mutant, showed any change in titanization capacity or endogenous superoxide accumulation relative to wild type. However, *sod2* Δ exhibited defects in the accumulation of superoxide (Fig 8B: right) and reduced formation of large cells (Fig 8B: left). *SOD2* encodes Mn-Sod2, the major superoxide dismutase in the mitochondrion, where it converts superoxide to H_2_O_2_ and is required for resistance to oxidative stress [61]. *SOD2* has been previously show to be required for growth at 37°C[61]. Examination of the cultures showed that *sod2* Δ mutant cells proliferate as yeast under titan inducing conditions (37°C, 5%CO_2_) (Fig 8C). While temperature has been identified as one of the conditions influencing robust titan cell induction *in vitro,* titanization can still be observed at 30°C, albeit at reduced levels [9, 11]. We therefore compared cell size in *SOD2* and *sod2Δ* cells at 30°C, 5%CO2, in titan-inducing medium (Fig 8D). There was no difference in growth between *SOD2* and *sod2Δ* cells 30°C (Fig 8D) Treatment with DPTA NoNoate increased the proportion of large, high ploidy cells for *SOD2* and there was a further increase for the *sod2Δ* mutant (Fig 8D). Together, this suggests that detoxification of mitochondrial superoxide by Sod2 is required to enable the yeast-to-titan switch under physiological conditions (37°C, 5%CO_2_).

**Figure 8:**
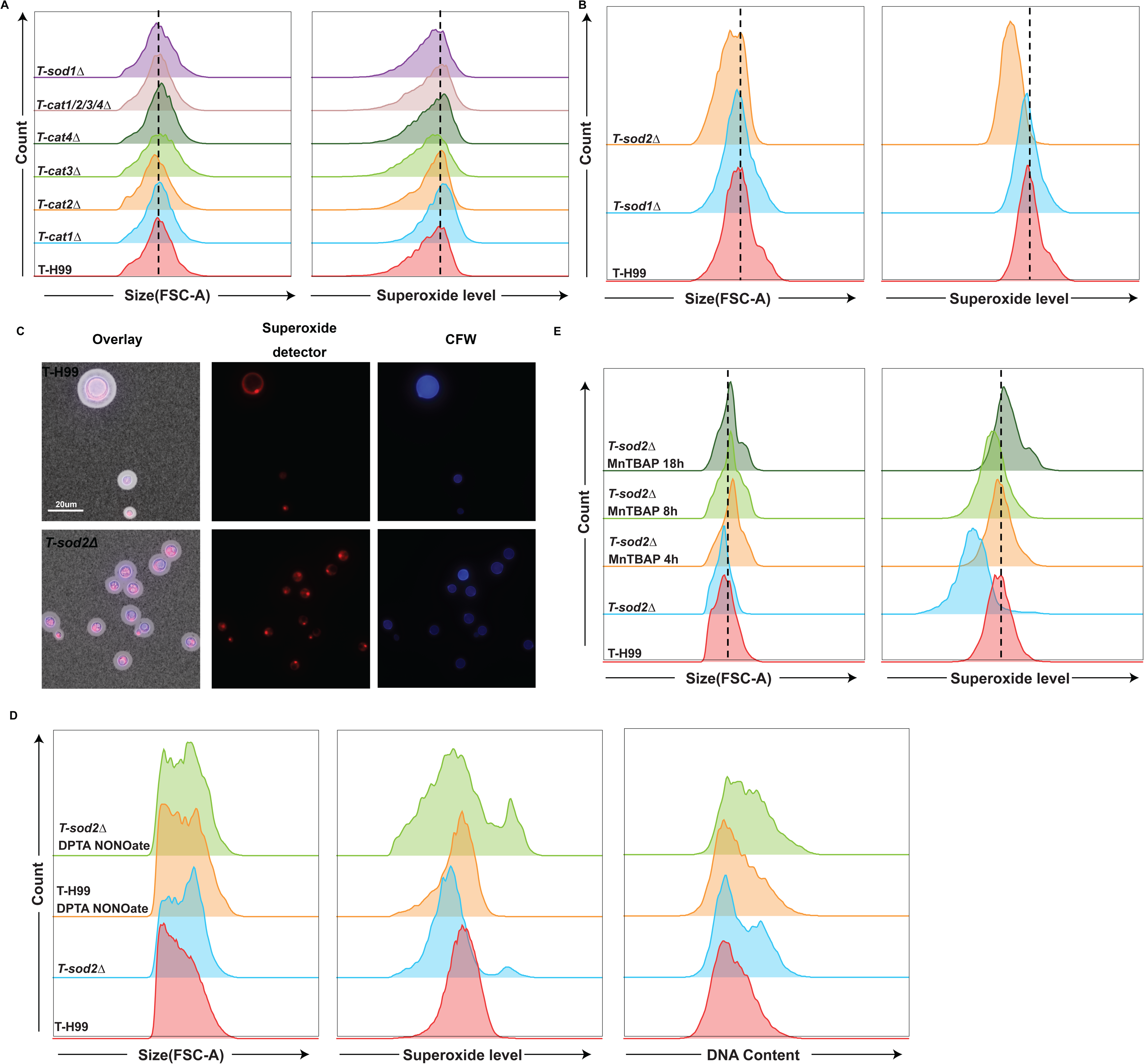
Superoxide Dismutase (Sod) but not Catalase (Cat) activity is required for titan cell formation. (A) Histograms of cell size and superoxide levels of different mutants: *sod1Δ, cat1Δ, cat2Δ, cat3Δ, cat4Δ* single mutants, and the cat1/2/3/4Δ quadruple mutant. All mutants and H99 were titan induced for 24hr and stained with superoxide detector, then analysed by flow cytometer for size (FSC-A) (*left*), and superoxide level (*right*). (B) Histograms of superoxide levels and cell size of *SOD* mutants: *sod1Δ* and *sod2Δ* with H99 as the control. All mutants and H99 were titan induced for 24hr and stained with superoxide detectors, then analysed by flow cytometer for size (FSC-A) (*left*), and superoxide level (*right*). (C) Microscopy images of *sod2⊗* growth after titan induction compared to H99. Cells were stained with superoxide detector (*red*), calcofluor white (CFW) for cell wall (*blue*) and India ink for capsule. Scale bar: 20 μm. (D) Histograms of cell size and superoxide levels of *sod2Δ,* and *sod2Δ* treated with 0.5mM MnTBAP for the initial 4h or 8h of titan induction and then washed away or for the entire 18hr titan induction period. <u>H99 incubated under similar conditions serves as the control</u>. All cells were stained with superoxide detector and then analysed by flow cytometer for size (FSC-A) (*left*), and superoxide level (*right*). (A&B&D) Single cells were identified by FSC-A vs FSC-H to remove doublets. Dotted lines indicate MFI of H99 titanized cells. “T” indicates pre-titanized

We hypothesized that, in titanizing cells at 37°C, the *sod2*Δ mutant accumulates high levels of superoxide in the mitochondria, resulting in failure to survive the yeast-to-titan transition. The dynamic nature of superoxide during titanization (Fig 4A) might indicate different requirements for superoxide detoxification over the course of induction. To test this, *sod2*Δ cells were placed under inducing conditions (37°C, 5%CO_2_) and treated with MnTBAP (1mM) for the initial 4hr or 8hr only, and then washed and returned to inducing conditions, or maintained in MnTBAP for 18hr. As shown in Fig 8E, MnTBAP treatment of 4 hrs was sufficient to reduce *sod2Δ* superoxide levels comparable with untreated *SOD2* titanized cells (Fig 8E). Treatment of *sod2*Δ cells for a further 8 or 18hr increased the overall proportion of large cells, yielding a population similar to *SOD2* titanized cells (Fig 8E). These data reveal that *SOD2* is required for initial detoxification for superoxide during titanization, as initial recovery of superoxide dismutase function is sufficient for *sod2* Δ to recover titanization defects.

## Titanized cells exhibit high nuclear superoxide and increased DNA damage

Microscopy of titanizing cells revealed that, during the later stages of titanization, superoxide becomes localised to the nucleus (Fig 9A). Zhao et al. recently showed that genotoxic stress leads to cryptococcal cell polyploidization [62]. We therefore hypothesize that this excessive endogenous superoxide within the nucleus is a source of genotoxic stress, and predicted that the accumulation of superoxide leads to DNA double-strand breaks (DSBs). To contend with this, cells must undergo DSBs repair through homologous recombination or non-homologous end joining [63]. We therefore measured the accumulation of Rad51, a member of the RecA-family that is a marker of DSBs [64–66], in titan and yeast-phase cultures. We used an anti-Rad51 antibody against whole cell lysates from titan-induced and yeast-phase cells, compared to UV-treated yeast cells as a positive control. In H99 titanized (T-H99) cultures, we detected a strong signal for Rad51, comparable with UV-treated cells, while only limited Rad51 was detected in yeast-phase cultures (Fig 9B). These data suggest that, in addition to high overall endogenous superoxide levels and mitochondrial polarization, superoxide in the nucleus may trigger the accumulation of DNA DSBs during titanization.

**Figure 9:**
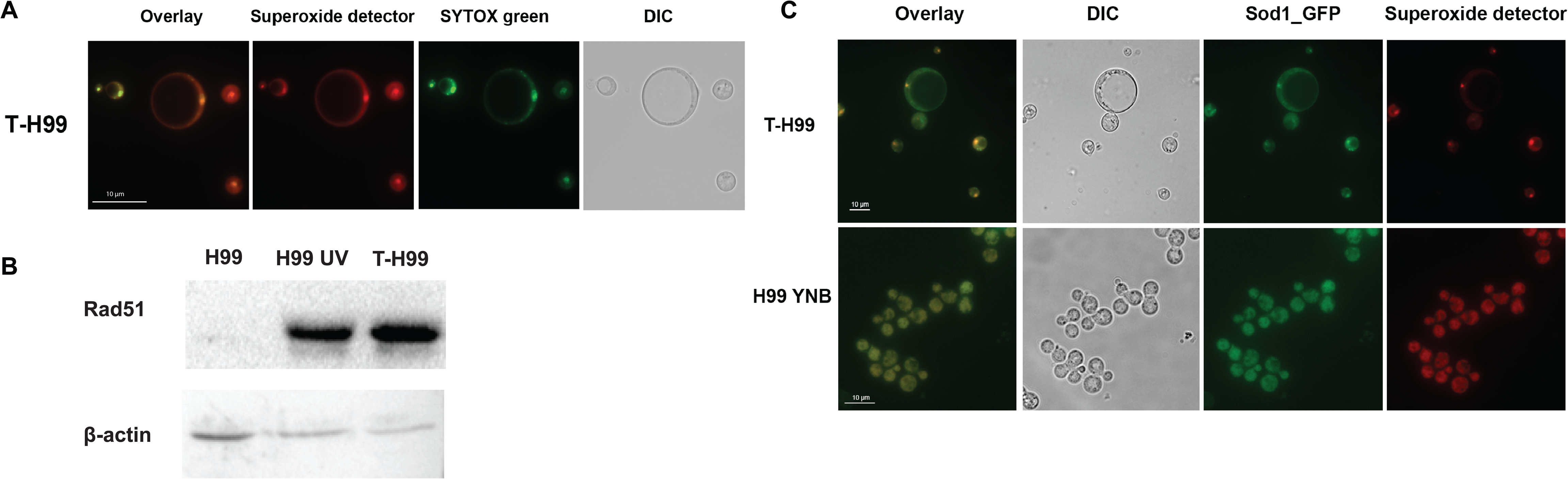
Sod1 and superoxide localise to the nucleus of yeast and titan cells under titan inducing conditions. (A) Superoxide co-localizes with nuclei in titanized cells. Fluorescent microscopy images of H99 titanized cells following staining with superoxide detector dye (*red*) and SYBR green for the nuclei (*green*). Scale bar: 10 μm. (B) Western blot for Rad51 protein levels in YPD-grown yeast-phase cells (H99), UV treated yeast cells (H99 UV) and titanized cells (T-H99, 24hr) with actin loading control. (C) CnSod1 subcellular localization at the start of titanization (H99 YNB) and 24hr post-induction (T-H99). Green: Sod1-GFP, Red: superoxide detector. Scale bar: 10 μm, “T” indicates pre-titanized.

## Sod1 is localised to the nucleus after titan induction

Our observation of nuclear accumulation of superoxide in titanized cells (Fig 9A) raised the question of how *C. neoformans* copes with oxidative stress. *C. neoformans* Cn/Zn-Sod1 is predicted to be mitochondrial and cytoplasmic, yet *S. cerevisiae* Cn/Zn-Sod1 has been shown to translocate into the nucleus upon exposure to high oxidative stress [67]. Therefore, we determined the subcellular localization of CnSod1 via N-terminal tagging with GFP under its native promoter. During yeast-phase growth (in YNB), GFP-Sod1 localises to the cytoplasm (Fig 9C). However, upon titan-induction, GFP-Sod1 was additionally observed in the nucleus, co-localised with superoxide (Fig 9C). This subcellular localization suggests that CnSod1 is important for regulating endogenous ROS levels in the nucleus during titanization. Despite this, loss of *SOD1* had no apparent impact on the capacity of yeast to convert into titan cells (Fig. 8A, B).

## *SOD1* regulates genomic integrity of titan daughters

Given that genotoxic stress can cause chromosomal rearrangement, and that titan cells often produce aneuploid daughters, we predict that the altered regulation of ROS in the nucleus after the deletion of *SOD1* might promote genomic instability in titan daughters [8, 62, 68]. To test this, we first induced titan cells in either *sod1* Δ or *SOD1* isolates for 48hr and then separated out daughter cells via passage through 4μm filters. This excludes the majority of yeast phase cells and enriches for titan daughter cells (2-3 μm). Daughter cells were plated on YPD at low concentration to isolate single cells. After 48 hr incubation at 30°C, for H99, all colonies were uniform in size. However, for *sod1* Δ two different colony sizes were observed: approximately 50% of *sod1* Δ derived colonies were smaller, indicating slower growth rate (Fig S5). Because aneuploidy is often correlated with growth defects, we compared the DNA content of cells from typical sized (B) and small (S) colonies of *sod1* Δ with H99-derived colonies as a control [69]. As shown in Fig 10A, *sod1* Δ titan daughters showed more variation in base DNA content compared with H99. In addition, slow growing *sod1* Δ *-*derived colonies exhibited a higher frequency of increased DNA content (aneuploidy) compared with either H99*-*derived or large *sod1* Δ-derived colonies (Fig 10A right panel). In order to validate the presence of aneuploidy, we also re-cultured these slow growing daughters in liquid YPD for 24hr and determined DNA content. As shown in Fig 10B, the increased DNA content for slow growing *sod1*Δ titan daughters was stable after re-culture. In summary, these data demonstrate that CnSod1 regulation of ROS in the nucleus is important for genome integrity during titanization and the production of euploid offspring.

**Figure 10:**
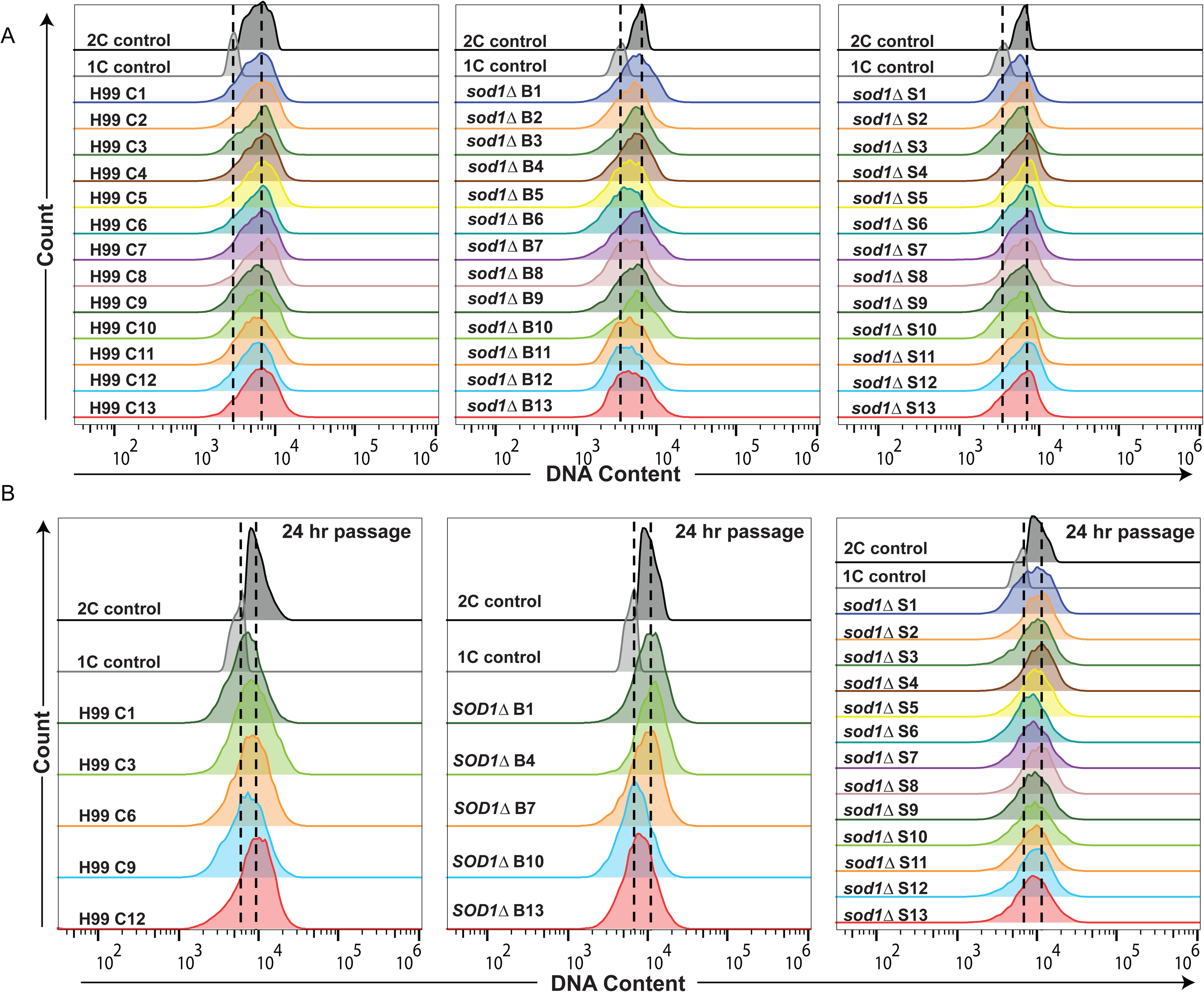
Loss of *SOD1* impacts titan daughter ploidy. H99 and *sod1Δ* cultures induced to form titan cells were filtered to enrich for cells <4 μm (titanides) and plated onto YPD plates. DNA content (DAPI) was analysed for colonies (A) after 48 hr directly from YPD plates and (B) after further 24 hr overnight YPD culture. (A&B) Histograms of DNA content, left): H99 colonies, middle): *sod1Δ* big colonies, right): *sod1Δ*small colonies. YPD grown yeast cells from matched parent strains and MFI 1C and 2C populations were used for 1C and 2C peaks. Dotted lines indicate the MFI of 1C and 2C peaks. *sod1Δ* B*: sod1Δ* -derived big colonies, *sod1Δ* S: *sod1Δ* - derived small colonies. T” indicates pre-titanized.

## Discussion

During infection, titan cells impair the host response directly and indirectly by blocking phagocytosis of small cells, resisting host stress, and influencing drug resistance [7, 8, 70, 71]. Here, we demonstrate that the host immune response also influences and contributes to titanization. Titan cells form in response to the host lung environment and can be generated via host-relevant stimuli *in vitro* [5, 6, 10–12].

Yet, despite strong evidence that *in vitro* stimuli are sufficient to induce titan cells, *in vitro-*generated titan cells are generally smaller than their *in vivo-*derived counterparts and produce diploid, rather than haploid or aneuploid, daughter cells [8–12]. Here, we observed *in vitro* that when pre-titanized cells were engulfed by macrophages, a proportion continued to undergo further cell enlargement within the phagolysosome. This phenomenon raised the hypothesis that host immune cell factors also contribute to titanization. By testing this hypothesis, we now are able to present a molecular model showing a role for endogenous superoxide in regulating the yeast-to-titan transition and also show how the host influences this process (Fig 11).

**Figure 11:**
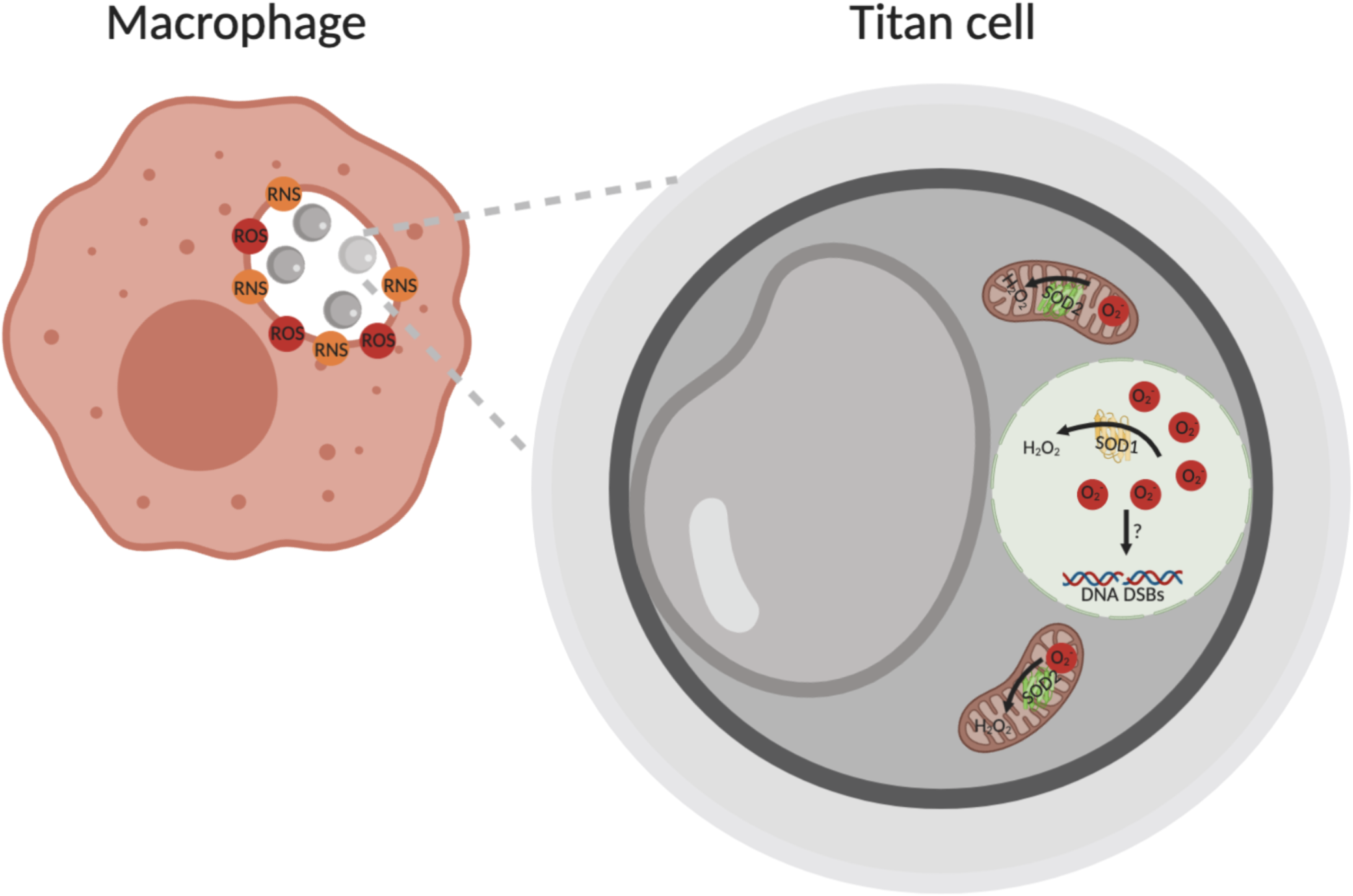
Summary of interaction between macrophages and *C. neoformans* titan cells. Macrophages bind and internalize cryptococcal cells through the formation of a phagosome, where they are exposed to antimicrobial effectors including ROS/RNS. However, for titan cells these host-derived oxidative and nitrosative stresses can lead to increases both in endogenous superoxide and cell size. This endogenous superoxide production is required for titan cells formation, and is detoxified by superoxide dismutases Cu/Fe-Sod1 and Mn-Sod2. Mn-Sod2 is a mitochondrial dismutase required for initial detoxification of superoxide during titanization. Cn/Fe-Sod1 translocates to the nucleus after titan induction, where it detoxifies genotoxic superoxide in the nucleus. Finally, this genotoxic superoxide is associated with DNA double strand breaks (DSBs), and loss of detoxification increases progeny aneuploidy.

First, by comparing cells exposed to either LPS+IFN-γ or IL-4 activated phagocytes, we found a positive correlation between increased titanization (cell size and ploidy) and M1 immune cell activation. We further demonstrated that host RNS, more than host ROS, contributes to the accumulation of endogenously generated superoxide within the fungal cell. By exploring the mechanism underlying these changes in endogenous ROS levels, we found that the fungal mitochondrion is highly activated by exogenous RNS, triggering increased production of superoxide, which is required for titanization. Blocking superoxide accumulation through the addition of a Sod mimetic drug blocks titanization, while addition of H_2_O_2_ or loss of catalase activity that leads to H_2_O_2_ accumulation has no impact on titanization. Together, these data indicate that superoxide, but not H_2_O_2_, mediates the yeast-to-titan transition (Fig 10).

Second, we show that even in the absence of external RNS, cryptococcal cells in titan inducing conditions exhibit increased endogenous superoxide, which has both mitochondrial and nuclear localization and is dynamic over time and across the population. Free ROS can damage macromolecules and membranes; however, they also represent crucial signalling molecules involved in fungal growth, differentiation and infection [42, 72]. The spatiotemporal regulation of ROS production is necessary for sexual development in *A. nidulans*, perithecia formation in both *Neurospra crassa* and *Podospora anseria,* and differentiation of virulence-relevant appressoria and penetration hyphae in *Magnaporthe oryzae* [39, 73–75]. Analogous to these other systems, we propose that, in *C. neoformans,* the spike in superoxide production controls the formation of titan cells that are crucial for both fungal survival and pathogenesis.

In *C. neoformans,* two Superoxide Dismutase (SOD) enzymes neutralize superoxide: Sod1 and Sod2. Both enzyme activities are required for resistance to oxidative stress, however they have different roles in pathogenesis [30, 61]. *SOD2* is required for high-temperature growth and targets superoxide generated by mitochondrial Complex III [30, 61]. During titanization, detoxification of mitochondrial superoxide by Sod2 is required for fungal cell growth and the initial switch to titan cells. Loss of *SOD2* reduces growth under titan inducing conditions and prevents cells from making the switch to titan morphology, suggesting that factors that perturb *SOD2* activity or otherwise influence mitochondrial ROS will also perturb titanization. Consistent with this, Trevijano-Contador et al. demonstrated that addition of the mitochondrial inhibitor sodium azide increased the size and frequency of titan-like cells in their *in vitro* assay [10].

Interestingly, previous work showed that deletion of *SOD1* in C*. neoformans* results in reduced virulence and increased sensitivity to iNOS-mediated macrophage killing [30, 61]. In addition, the *yap1Δ* mutant exhibits reduced dissemination to the brain and slightly reduced overall virulence [59]. We demonstrate that the *yap1Δ* mutant fails to produce high ploidy titan cells and has a defect in titanide production, and that loss of *YAP1* impairs regulation of *SOD1.* Although *SOD1* is not required for titan cell formation, detoxification of nuclear superoxide by Sod1 is required for the generation of haploid progeny in the face activation of DNA Double Strand Break repair (Fig 9 and 10). Specifically, we demonstrate that Sod1 subcellular localization changes over time, translocating to nuclei in induced cells. This localization pattern suggests that Sod1 is involved in the regulation of ROS accumulation in the nucleus, and analysis of the *sod1Δ* mutant suggests that altered regulation may affect genome integrity of titan offspring, leading to the generation of aneuploid progeny.

Finally, we detected a strong signal for the DSB repair protein Rad51 in induced cells, indicating that there is high DNA damage accumulated during the yeast-to-titan switch. A recent paper reported that *Cryptococcal* polyploidization during infection is induced by unknown genotoxic stresses causing DSB [62, 68]. Combined with this, we predict high DNA damage is caused by accumulation of endogenous superoxide in the nucleus. In the future, the relationship between DSBs and titanization should be further investigated. Further research will also study the molecular mechanisms by which the superoxide signal is transduced, resulting in titan cell formation.

Together, our data link previous work on the role of major host response molecules to controlling *C. neoformans* proliferation and the morphological switch from yeast-phase growth to titan phase growth, a major mechanism by which *C. neoformans* evades the host. These findings pave the way for future work investigating the underlying regulation of fungal morphogenesis by host-derived and endogenously generated reactive species, including possible chemotherapeutic strategies to block the yeast-to-titan switch.

## Acknowledgements

We are grateful to the University of Birmingham Biomedical Services Unit as well as the doctors, patients, and their families involved in the ATCA Lusaka trial. We are grateful to the Hiten Madhani lab for their creation of the 2016 *Cryptococcus* deletion collection, from which the *sod2Δ* mutant was derived (NIH funding (R01AI100272)) and to the Fungal Genetics Stock Center for strain upkeep. We additionally acknowledge publically available resources made available through FungiDB, funded by US National Institute of Allergy and Infectious Diseases (Contract HHSN75N93019C00077), with additional support from the Wellcome Trust (Resource Grants 212929 & 218288).

## Materials and methods

### Strains, media and growth conditions

*C. neoformans* H99 was kindly provided by Andrew Alspaugh, Duke University, NC, USA. Zc1, Zc8 and Zc12 clinical isolated were collected from ACTA Lusaka Trial in Lusaka, Zambia [76, 77] and kindly provided by Tihana Bicanic and Matthew Fisher. Strains were cultured routinely on YPD (1% yeast extract, 2% bacto-peptone, 2% glucose, 2% bacto-agar) plates stored at 4°C. Titan cell *in vitro* induction was performed as described in [11].Cells were pre-cultured at 37°C, 150rpm, in 10ml YNB without amino acids (Sigma-Aldrich) in which 2% of glucose was added according to manufacturer’s protocol. After 24 hrs, cells were normalized to OD_600_=0.001 in 1x PBS (phosphate-buffered saline) with 10% of Heat inactivated Fetal Bovine Serum (HI-FBS) (Sigma-Aldrich F9665) and incubated at 37°C, 5%CO_2_ for 24 hrs unless otherwise indicated.

## Strain list

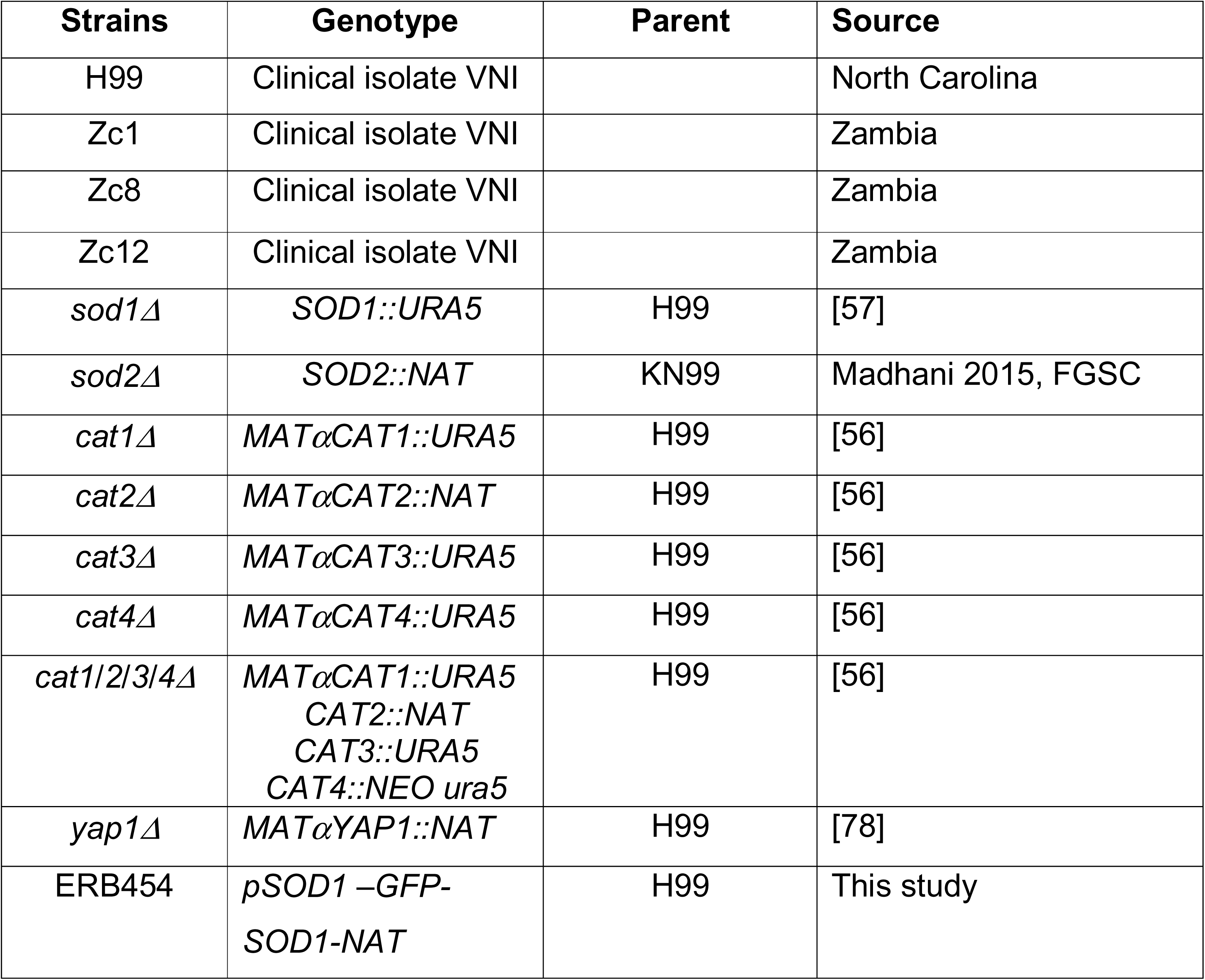

Strain list:
| Strains | Genotype | Parent | Source |
| --- | --- | --- | --- |
| H99 | Clinical isolate VNI |  | North Carolina |
| Zc1 | Clinical isolate VNI |  | Zambia |
| Zc8 | Clinical isolate VNI |  | Zambia |
| Zc12 | Clinical isolate VNI |  | Zambia |
| *sod1Δ* | *SOD1::URA5* | H99 | [57] |
| *sod2Δ* | *SOD2::NAT* | KN99 | Madhani 2015, FGSC |
| *cat1Δ* | *MATαCAT1::URA5* | H99 | [56] |
| *cat2Δ* | *MATαCAT2::NAT* | H99 | [56] |
| *cat3Δ* | *MATαCAT3::URA5* | H99 | [56] |
| *cat4Δ* | *MATαCAT4::URA5* | H99 | [56] |
| *cat1/2/3/4Δ* | *MATαCAT1::URA5*<br>*CAT2::NAT*<br>*CAT3::URA5*<br>*CAT4::NEO ura5* | H99 | [56] |
| *yap1Δ* | *MATαYAP1::NAT* | H99 | [78] |
| ERB454 | *pSOD1 –GFP-*<br>*SOD1-NAT* | H99 | This study |

### Generation of Sod1-GFP

*C. neoformans* strain ERB454 expressing GFP-Sod1 via the native *SOD1* promoter with Neurceothrycin (NAT) resistance was generated as follows: The eGFP sequence was amplified from pHGNAT plasmid (pCN19, a kind gift from Connie Nichols and Andy Alspaugh) using primers ERB574 and ERB575 [79]. The 600bp promoter of *CAT3* (CNAG_00575) (pCAT3) was amplified using primers ERB524 (ccggaagcttCTCACCTTTCGCATTCCTTT) and ERB525 (TCGCCCTTGCTCACCATAGTGCGTCATAATTACCGTGG) to introduce HinDIII and BamHI flanking restriction sites and an internal NdeI restriction site. Primer ERB525 additionally enabled overlap PCR with the eGFP fragment. The pHGNAT plasmid was modified to excise the *HIS3* promoter sequence and eGFP sequences via HinDIII and BamHI restriction digest. The linearized plasmid backbone was ligated with the p*CAT3*-GFP PCR construct, resulting in plasmid pCAT3GNAT (pEB048) enabling excision of the *CAT3* promoter. The *SOD1* promoter and 5’UTR (CNAG_01019) were amplified using primers ERB532 (ccggaagcttCACACTCGGCACTATCGG) and ERB533 (ggcatatgAGGAAGATATGGATGAGAGGT) (225bp product) The p*CAT3* insert in pEB048 was excised by HinDIII/NdeI digest and replaced with the SOD1 promoter upstream of eGFP to produce plasmid pSOD1GFP-NAT (pEB060). A 1.1kb region encoding the *SOD1* ORF (CNAG_01019) was amplified using primers ERB585 (ggatccATGGTCAAGGTACGTACAGTA) and ERB586 (cctgaggATGCAATGTTGGAATGTACT) to incorporate BamHI and SauI restriction sites in frame. The resulting construct was cloned via BamHI and SauI into the plasmid pEB060 in-frame with eGFP to produce plasmid pEB061 (pSOD1-GFP-SOD1-NAT). Genomic integration into H99 was performed using the biolistic transformation method as previously described[80, 81].

#### Oxidative stress plating

*C. neoformans* cells were collected from YNB overnight (0hr) and 18hrs post titan induction cultures, then plated in 10-fold serial dilutions from 1 × 10^6^ cells/ml onto YNB or YNB plus 0.5mM H_2_O_2_.Plates were incubated at 30°C for 2 days and imaged.

#### Cell sizes measurement

Cell diameter was measured using FIJI, with frames randomly selected, all cells in a given frame analysed and at least 5 images acquired per sample for each of two independent runs. Total cell number of each samples was >200. All statistical analyses were performed using Graphpad PrismV8.

#### Superoxide staining

Cells were collected and washed twice with 1 x PBS, then stained with superoxide detection reagent (Enzo Life Sciences) (1:500 dilution). Live stained cells were imaged with Zeiss Axio Observer and analysed for fluorescent intensities with Attune Nxt Flow cytometer YL2 channel.

#### JC-1 staining

The Abcam JC1-mitochondrial membrane potential assay kit was used according to the manufacturer’s instruction. Live stained cells were imaged with Zeiss Axio Observer and 10,000 cells analysed for fluorescent intensities with Attune Nxt Flow cytometer BL1 and BL2 channels. Doublets and clumps were excluded using recommend gating system of SSC-H vs SSC-W followed by FSC-H vs. FSC-W, and auto-fluorescence was excluded using unstained cells. Gates for 1C and 2C were established with H99 haploid controls cultured in YPD liquid media.

#### Flow cytometry for cell ploidy

Cells were fixed and stained according to the protocol of Dambuza et al., [11]. Cells were fixed with 50% methanol for 15mins, washed 3 times with 1 x PBS and stained with 0.3ug/ml DAPI. Cells were analysed for DNA content using an Attune NxT Flow cytometer in the VL-1 channel. All flow cytometry data were analysed with FlowJo. Doublets and clumps were excluded using recommend gating system of FSC-A vs. FSC-H, and auto-fluorescence was excluded using unstained cells (S2B Fig). Gates for 1C and 2C were established with H99 haploid controls cultured in YPD liquid media.

#### DPTA NONOate treatment

After culturing in YNB overnight, cells were induced for titanization as described above with adding extra 0.3uM of DPTA NONOate (Enzo Life Sciences) which was resuspended in DMSO (Dimethyl sulfoxide). 48h post-induction, cells were analysed with Attune Nxt Flow cytometer both for cell sizes and superoxide production with untreated cells as a control.

#### Glucose oxidase treatment

After culturing in YNB overnight, cells were induced for titanization as described above with adding extra 1mU of glucose oxidase (generates 1nmoles H2O2/min). 24h post-induction, cells were analysed with Attune Nxt Flow cytometer both for cell sizes and superoxide production with untreated cell as a control.

#### MnTBAP treatment

After culturing in YNB overnight, cells were induced for titanization as described above with adding extra 1mM MnTBAP which was resuspended in PBS plus NaOH (PH=7.0). After 24h incubation in dark, cells were analysed with Attune Nxt Flow cytometer both for cell sizes and superoxide production with untreated cell as a control. For *sod2Δ* treatment, 500uM MnTBAP was added into for initial 4h or 8h, then drug was washed away with PBS and cells were resuspended with 1 x PBS with 10% of HI-FBS (titan-induction buffer).

#### Western blotting

*C. neoformans* cells were grown in titan induction condition for 48-72h and YPD overnight. For overnight YPD cells, they were divided into two groups: 1) YPD-grown cells used as a negative control and 2) cells were treated with UV 100 J/m^2^ for 1h used for a positive control. Cells were collected by centrifugation and washed two times with cold PBS, and cell pellets were flash frozen with liquid nitrogen (store at -80°C). Cell pellets were suspended with 2ml lysing buffer (50 mM L Tris-HCl [pH 7], 150mM NaCl, 50mM Tris, 5mM EDTA and 1% Triton X-100) and 0.5ml of o.5-mm glass beads (Millipore Sigma) on ice. After that, the cells were broken using a bead beater for 5 cycles of 2x30 s at the maximum speed with 1Lmin cooling on ice. Broken cells were centrifuged at maximum speed at 4°C for 10min, then supernatant was transferred into a sterile Eppendorf tube. Total protein was diluted with 1xPBS and the concentration was determined using a Bradford reagent (Sigma-Aldrich). Samples of 50 µg of protein were separated on a 10% SDS PAGE gel, then transferred to a nitrocellulose membrane. The blot was blocked in 10% BSA in 1 x PBS with 0.1% Tween 20. Both Rad51-specific and anti-actin antibodies were used at a 1:2,000 dilution, and the blot was incubated overnight at 4°C. HRP-conjugated anti-rabbit IgG secondary antibody was used at a 1:4,000 dilution, and the blot was incubated for 2 Lh at 25°C. Finally, the blot was incubated with Clarity Western ECL substrate (Bio-Rad) according to the manufacture’s instruction, and the image was acquired through a ChemiDoc Touch Imaging system (Bio-Rad).

#### J774.1 cell culture

For most *in vitro* infection assays, we used the J774.1 murine macrophage cell line. Cells were routinely passaged in Dulbecco’s modifies Eagle medium (DMEM+) culture media with serum (DMEM, low glucose, from Sigma-Aldrich; 10% HI-FBS; 1% penicillin-streptomycin solution from Sigma-Aldrich; 1% 200mM L-glutamine from Sigma-Aldrich) and 100 ng/ml LPS (Sigma) and 1000 U/ml IFN-γ (R&D Systems) were added as indicated overnight. All assays were performed with cells in passages 4 to 10. 24h before infection, 1x 10^5^macrophages J774.1 were seeded into 12-well plastic plates in 1 ml DMEM culture media with serum (DMEM+) and activated with 10 U/ml IFNγ.

#### *In vitro* infection assays

Before infection, *C. neoformans* cells were collected from titanization culture and washed three times with sterile 1 x PBS. After normalising to 1 10^6^ cells/ml in 300 μl DMEM+, cells were opsonized with 10μg/ml anticapsular x 18B7 antibody by incubating at 37°C for 30min. Then opsonized *C. neoformans* cells were added into activated macrophages or monocytes to give a multiplicity of infection (MOI) ratio of 10:1. After 2 hrs (time point 0), infected wells were washed at least three time with pre-warm 1 x PBS until all un-engulfed *C. neoformans* cells were removed from the wells. Infection assays stopped after another 22h, monocytes and BMDMs were lysed with ice-cold water and *C. neoformans* were recovered for further analysis.

#### J774.1 macrophages live imaging assay

At time point 0 of phagocytosis assays, extracellular cells were removed by washing with warm fresh DMEM+. All samples were maintained at 37°C, 5%CO_2_ in the Nikon Ti-E microscope chamber and imaged for 18h via time-lapse. Images were taken every 15 min and compiled into single movie files for analysis using NIS elements or FIJI software. Movies then were monitored visually for cell size changes over time.

#### Flow cytometry for cell size and superoxide level after *in vitro* infection

Following co-culture, macrophages were lysed with ice-cold water and both fungi and macrophages were collected, centrifuged, and washed once with 1xPBS. Cells were stained with anti-IgM (Crp127) FITC(BL1-A) and superoxide detector dye (Enzo Life Sciences) [82]. Cells including both fungal and macrophages were analysed for using an Attune NxT Flow cytometer. Doublets and clumps were excluded by gating on FSC-A vs FSC-H, and fungi were identified by positive capsule staining (BL1-A). BL1-A+ cells were then examined for size (FSC-A) and superoxide staining (YL2-A).

#### Human primary monocytes isolation and culture

All human tissue work was performed as approved by the University of Birmingham Ethics Committee under reference ERN_15-0804b. On day of draw, 25-30 ml of blood was withdrawn from anonymized healthy volunteers by venepuncture and was directly used for monocytes isolation with double layer percoll. Before adding the blood, two of different densities of percoll were layered in the universal tubes with 6 ml of 1.079 on the top and 6 ml of 1.089 at the bottom. After 6 ml of whole blood was layered over dual gradient, the universal tubes were centrifuged at 150g for 8mins, followed by 10mins at 129g without acceleration or breaks. The white disc of peripheral blood mononuclear cells (PBMCs) was removed from the top and mixed with red blood cell lysis buffer to clear away red blood cells. After centrifuge for 6 mins at 400g, PBMCs were washed twice in cold 1 x PBS (centrifuged at 1000 g for 6 mins in between), resuspended in warm RPMI 1640 with 5% HI-AB serum, counted, and then plate onto 12 well plate at 1 x 10^6^ cells per well. The covered plate was incubated at 37°C, 5%CO_2_ overnight, after which non-adhere cells were removed by washing with fresh RPMI.

**Mice** Male B6.129S-*Cybb^tm1Din^*/J or C57BL/6 mice were used at 10-33 weeks of age and were maintained in individually-ventilated cages under specific pathogen-free conditions at the Biomedical Research facility at the University of Birmingham (Birmingham, UK). Animals were provided with food and water *ad libitum.* All experimentation conformed to the terms and conditions of United Kingdom Home Office license for research on animals (PPL/PBE275C33) and the University of Birmingham ethical committee.

#### Murine bone marrow derived macrophages

For each experiment, rear legs from one mouse were disinfected for 1 minute in 70% ethanol. The muscles were removed and both extremities of the femurs and tibias were cut to flush the bone marrow out with complete medium (RPMI 1640 with L-glutamin, 0.1 mg/mL Penicilin/Streptomycin mix, 10% Heat-Inactivated Fetal Bovine Serum) into a 6 well plate using a small needle associated to a syringe. The bone marrows were put in suspension in 8mL complete media solution by gentle up and down with a p1000 pipette. These cell suspensions were filtered on a 100um mesh, transferred in a 50mL tube and the well rinsed with 2mL of complete media that were added to the cell suspension while rinsing the mesh. After a 3 minutes centrifugation at 400g, the cell pellets were resuspended in 10mL of complete media. Small aliquots of these cell suspensions were diluted at 1/10 before proceeding to leucocyte counting on a Neubauer Chamber under the microscope. Further to that count, the cells were centrifuged and resuspended in complete media containing 20ng/mL of mouse recombinant murine M-CSF in order to dispense 5x10^5^ cells per well of a 24 well plate in a final volume of 500uL (“day 0”). At day 3, 500uL of M-CSF complete media were added to the wells. At day 6 or 7, 500uL of culture media was removed and 500uL of new M-CSF complete media were added to the wells. The following day the wells were gently washed two to three time with warm complete medium in order to remove non-adherent cells and were incubated with complete medium supplemented or not with 10 ng/mL of recombinant murine IL-4 (Peprotech) or a mixture of 100 ng/ml LPS (Sigma) and 1000 U/ml IFN-γ (R&D Systems). The presence of adherent macrophages was confirmed by observation under microscope.

In some cases, bone marrow was frozen for future use. Following filtration through 100 um mesh, bone marrow cells were centrifuged, resuspended in 1mL of freezing media (L-glutamin supplemented RPMI 1640 containing 0.05 mg/mL Penicilin/Streptomycin, 45% Heat-Inactivated Fetal Bovine Serum, 10% DMSO), and transferred in a micro-vial to be frozen in a Mr Frosty cell container at -70°C overnight before being transferred in liquid nitrogen. Cells were recovered by bringing the micro-vial to 37°C in a water bath and slowly diluting with 9 mL warm complete media. After centrifugation, cells were resuspended in 10 mL complete media, small aliquots were diluted at 1/10 in a 0.2% trypan blue solution to identify dead cells, and live leucocytes were counted on a Neubauer Chamber under the microscope.

Further to this count, the cells were cultured as described above.

---

**Fig S1: Time-lapse movie of interaction between macrophages and *C. neoformans* cells. (**A) 18hr time lapse imaging of interaction between macrophages and H99 titanized or (B) YNB-grown cells.

**Figure S2:(A) Fungal cell size change was not affected by IL-4 stimulation.** *C. neoformans* pre-titanized cell size changes after incubation in RPMI in presence or absence of BMDMs, IL-4. *C. neoformans* cell size was assessed for individual cells (cell body diameter, excluding capsule; n>200). Data were assessed for normality by Shapiro-Wilk and analysed by one-way ANOVA, \*\*\**p*=0.0009, \*\**p*=0.0049. **(B)Flow cytometric gating for identification of fungal cells.** Single cells were first identified by FSC-A vs FSC-H, then fungal cells were gated by FSC-A vs BL1-A (capsule staining).

**Fig S3: JC-1 staining within titanized cells.** Representative micrographs for JC-1 staining of H99 titanized cells, green: JC-1 green fluorescence, red: JC-1 red fluorescence. Scale bar: 10μm T” indicates pre-titanized

**Fig S4: Microscopy images of cells treated with MnTBAP during titanization**. (A) Cells were stained with india ink and superoxide detector, Top panel: untreated cells, bottom panel: 1mM MnTBAP treated cells, red: superoxide detectors, blue: CFW. Scale bar: 20μm. (B) H99 YPD grown yeast cells were gated for singlets and MFI of 1C and 2C populations were used to identify 1C and 2C peaks. T” indicates pre-titanized

**Fig S5: H99 and *sod1****Δ***titan daughters re-culture plates.** *sod1Δ* titan daughters (centre and right) showed two different colony sizes on re-culture plates. Black arrows point small colonies.

