## Supplementary figures and images for "Host-derived Reactive Nitrogen Species mediate the *Cryptococcus neoformans* yeast-to-titan switch via fungal-derived superoxide"

### Supplemental Figure S2

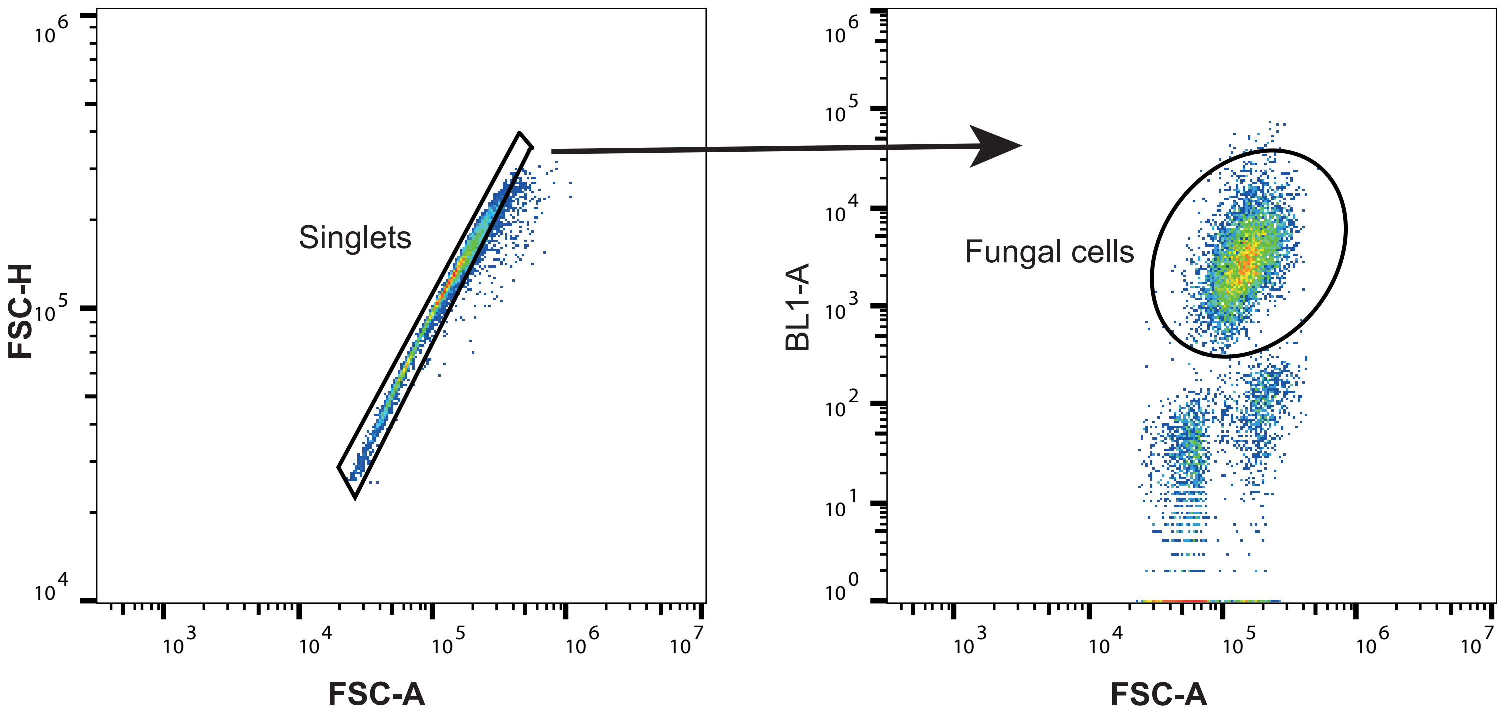

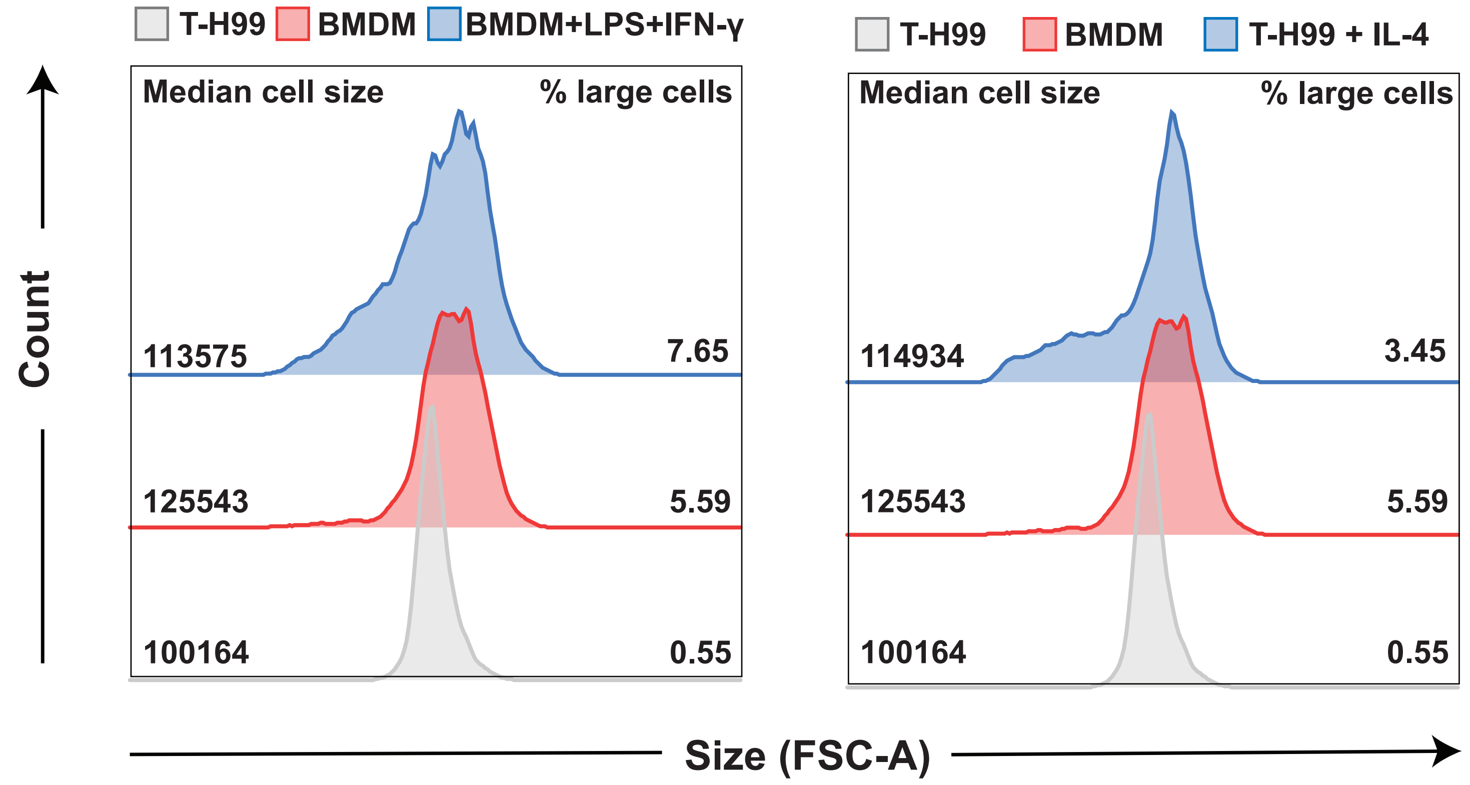

### Supplemental Figure S3

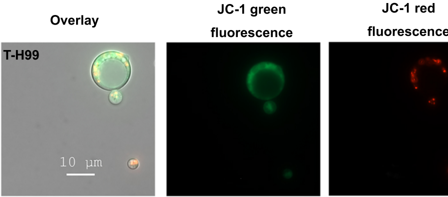

### Supplemental Figure S4

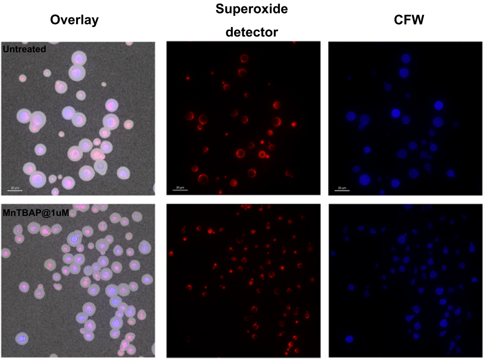

### Supplemental Figure S5

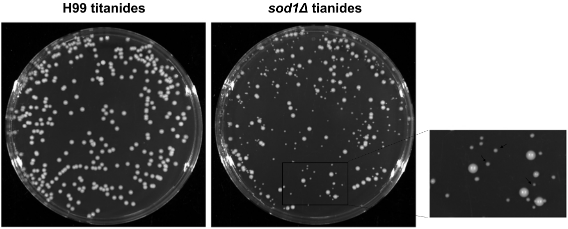
